# Infiltrative classical monocyte-derived and *SPP1* lipid-associated macrophages mediate inflammation and fibrosis in ANCA-associated glomerulonephritis

**DOI:** 10.1101/2024.03.13.584866

**Authors:** Yosta Vegting, Aldo Jongejan, Annette E Neele, Nike Claessen, Gal Sela, Koen H M Prange, Jesper Kers, Joris J T H Roelofs, Joost W van der Heijden, Onno J de Boer, Ester B M Remmerswaal, Liffert Vogt, Frederike J Bemelman, Menno P J de Winther, Perry D Moerland, Marc L Hilhorst

## Abstract

**Background:** Kidney macrophage infiltration is a histological hallmark of vasculitic lesions and is strongly linked to disease activity in anti-neutrophil cytoplasmic antibodies (ANCA)-associated glomerulonephritis (AGN). The precise mechanisms by which kidney macrophages influence local inflammation and long-term damage remain largely unknown.

**Methods:** Here, we investigate kidney macrophage diversity using single-cell transcriptome analysis of 25,485 freshly retrieved unfrozen, high-quality kidney CD45^+^ immune cells from five AGN patients, a lupus nephritis and nephrectomy control. Detailed subclustering of myeloid cells was performed to identify disease-specific macrophage subtypes. Next, transcriptome differences between macrophage subsets and disease serotypes were assessed. Findings were validated by immunostainings of an extended cohort of kidney biopsies and flow cytometric analysis of peripheral blood monocytes.

**Results:** Four main macrophage subsets were identified, including a classical monocyte-derived macrophage (MDM) subset expressing a chemotactic (*CXCL2, CXCL3, CXCL8*, *CCL3*) and pro-inflammatory (*IL1β, TNF*) set of markers and a osteopontin/SPP1^+^ lipid-associated macrophage (*SPP1* LAMs) subtype exhibiting distinctive upregulation of fibrotic genesets. AGN samples revealed a markedly increased proportion of CD163^+^ macrophages, predominantly composed of classical MDMs, accompanied by resident-like *C1Q* macrophages, and *SPP1* LAMs. An analogous trend was observed in the expansion of peripheral blood classical monocytes during active disease. The proteinase 3 (PR3)-AGN subtype exhibited heightened classical MDM infiltration and markers of acute inflammation, while interferon signaling and markers of chronicity were reduced compared to myeloperoxidase (MPO)-AGN.

**Conclusions:** Our findings highlight the expression of inflammatory and fibrotic genes by kidney macrophage subsets in AGN. Classical monocyte dysregulation might contribute to inflammation in the pathogenesis of AGN. Targeting these specific monocyte/macrophage subsets may potentially control the inflammatory cascade and attenuate resulting fibrosis in AGN and kidney disease in general.

**Key points:**

- Classical monocyte-derived macrophages are predominant in ANCA-associated glomerulonephritis and exhibit chemotactic and pro-inflammatory markers
- Osteopontin/SPP1^+^ lipid-associated macrophages (*SPP1* LAMs) show distinctive upregulation of fibrotic genesets
- Understanding of the macrophage immune response supports exploration of macrophage-directed therapies for the treatment of autoimmune kidney diseases

## Introduction

Anti-neutrophil cytoplasmic antibodies (ANCA)-associated vasculitides comprise a group of necrotizing small vessel vasculitides associated with autoantibodies against granular enzymes, proteinase-3 (PR3-ANCA) and myeloperoxidase (MPO-ANCA), located in monocytes and neutrophils^1^. While there are clinical and histological distinctions among these two serological subtypes, kidney involvement, termed ANCA-associated glomerulonephritis (AGN), is frequent in both, and characterized by severe glomerular inflammation without evidence of immune complex deposition^2-4^. Despite better treatments, high risks of renal disease, side effects, and relapses persist. Unraveling immune mechanisms in AGN is crucial for developing early and effective management strategies.

Kidney macrophage infiltration is a key histological feature of AGN^5^, but its function in the orchestration or resolution of local inflammation remains unknown. Current knowledge relies on immunohistochemistry studies, classifying macrophages according to the expression of a limited number of pro- or anti-inflammatory markers^6^. Recent insights show macrophages’ adaptable plasticity^9^, underscoring the necessity for deeper understanding of macrophage dynamics. Researchers have recently created multi-omic single-cell kidney cell atlases for healthy and diseased kidneys^10-12^, however, these atlases often lack detailed coverage and subclustering of macrophages. Furthermore, preservation methods for RNA sequencing vary in maintaining monocyte-derived macrophage integrity^13^ and no atlas specifically for AGN is available.

Here, we provide the first single-cell immune dataset of never-frozen, immediately processed kidney biopsies from patients with AGN. We performed detailed subclustering of myeloid cells and observed clear differences in kidney macrophage subtype distribution between AGN patients and a nephrectomy control. Inflammatory classical monocyte-derived macrophages (MDMs) were the most abundant macrophage subtype in AGN kidneys. We identified and validated a *SPP1* macrophage subtype expressing a highly specific set of markers associated with lipid metabolism and fibrosis. Our findings provide unique insights in the kidney macrophage landscape and supports the exploration of new therapeutic avenues in the treatment of autoimmune kidney diseases.

## Methods

Detailed information can be found in the Supplementary Methods.

### Study subjects and kidney tissue acquisition

Kidney tissue from AGN patients (n=5) and a lupus nephritis (LN) patient (n=1) was acquired during a biopsy procedure for a clinical indication. Healthy kidney tissue (n=1) was obtained from a kidney that was surgically removed due to a (non-invasive) papillary urothelial carcinoma (**Table S1-2**). The samples are referred to as AGN, LN and NC, respectively.

### Kidney tissue processing and flow cytometric cell sorting

Immediately after collection, kidney tissue was processed into a single-cell suspension using a modified protocol described earlier^14^, involving the use of digestion medium and filtering steps. Standard density gradient centrifugation was used in the NC sample to decrease sorting time according to manufacturer’s protocol. Kidney cells were incubated with a 4-color flow cytometry panel (**Table S3**) to sort CD45^+^ immune cells (**Fig. S1**) and estimate T cell and monocyte/macrophage proportions (**Table S4**). Surface protein abundance was determined in two samples (AGN3, NC) using an oligo-antibody TotalSeq-C cocktail [Biolegend] containing 130 unique cell surface antigens.

### Loading of cells and single-cell library preparation

To prevent alterations in transcriptomic profiles and cell integrity due to preservation methods, we implemented immediate processing. Therefore, after cell sorting or labeling, cells were loaded onto a Chromium chip [10x Genomics]. T and B cell receptor repertoires were determined in AGN3, NC. TCRαβ and BCR libraries were prepared with the V(D)J enrichment Kit v1.1 [10x Genomics]. cDNA synthesis, amplification, and sequencing libraries were made using either the single-cell 3’ Reagent v3.1 [10x Genomics] or the 5’ Reagent v1.1 [10x Genomics] (**Table S4**). Sequencing was performed on an Illumina NovaSeq 6000 platform at a target depth of 30,000 reads per cell.

### Single-cell data pre-processing and analysis

In short, after pre-processing and integration, we performed graph-based clustering and annotated the integrated dataset by mapping it onto the mature immune kidney cell atlas of Stewart *et al.*^15^ as reference (**Fig. S2a-c**). Clusters containing ≥ 5% of all myeloid cells were selected (**Table S5**). The resulting integrated myeloid-enriched dataset contained 4,194 cells, which were used as the basis for all analyses below.

We renormalized each sample separately, performed integration, clustered the integrated myeloid-enriched dataset at resolutions 0.3 and 1, and determined cluster marker genes and proteins for manual cell type annotation.

Genes differentially expressed between different conditions for cells of the same type were determined using a pseudobulk-based analysis^16^. The Benjamini-Hochberg false discovery rate (FDR) was used for multiple testing correction of the resulting *P*-values (significance; adjusted *P*-value <0.1). Geneset enrichment analysis (GSEA) was performed with selected geneset collections (Hallmark and the BioCarta, KEGG and Reactome subsets of the C2 collection) from the Molecular Signatures Database (MSigDB; v2023.1.Hs) with Benjamini-Hochberg FDR multiple testing correction (significance; adjusted *P*-value <0.1). Myeloid-enriched cells were scored based on a curated list of genes that regulate extra-cellular matrix (ECM) production in kidney tissue. Differences in cell type proportions between conditions were assessed using *propeller*^17^ with Benjamini-Hochberg FDR multiple testing correction (significance; adjusted *P*-value <0.1).

### Flowcytometric analysis monocytes

Flowcytometric analysis of cryopreserved peripheral blood monocytes was performed of 10 active AGN patients, 5 follow-up samples during stable disease and 5 age-and-sex matched healthy controls. PBMCs were incubated with a fixable viability dye eFluoro506 and 6 panels with surface antibodies (**Table S3**), and measured on a FACSCanto [BD]. Gating strategy is shown in **Fig. S1**. The mean percentage of monocyte subsets was calculated for each participant. Differences in monocyte proportions between conditions were assessed using *propeller*^17^ using moderated t-tests (either paired [active versus remission] or unpaired [rest]), Benjamini-Hochberg FDR corrected.

### Multi-color immunofluorescence stainings

*SPP1/ TREM2/PLIN2* LAMs and *S100A9* classical MDMs were detected by immunofluorescence stainings of an extended cohort of 36 kidney biopsies (22 AGN, 6 LN, 8 NC). Primary antibodies (**Table S3**) were stained overnight, followed by a stepwise incubation of the secondary antibodies to avoid cross-binding of goat antibodies. Whole-biopsy multi-color images were taken [Thunder Microscope system, Leica], maintaining consistent settings for all acquisitions. To visualize fibrosis, we conducted Picro Sirius Red (PSR) staining over the immunostaining in a subset of the biopsies (n=6). Total CD163^+^ macrophage area and the presence of S100A9/CD163 classical MDMs were semi-quantitatively analyzed with Fiji/ImageJ2 ^18^. CD163 and S100A9 thresholds were maintained consistent within each batch.

### Histopathological/clinical features

Paraffin sections of all kidney biopsies were subjected to a detailed histopathological examination by experienced nephropathologists (JR and JK). A chronicity score was calculated using the modified NIH chronicity index^19^, and a histological disease activity index was calculated based on cellular and fibrocellular crescents, tubulointerstitial nephritis (TIN), fibrinoid necrosis (FN) and Bowman’s capsule (BC) break. eGFR was calculated using the CKD-EPI 2021 formula^20^. Spearman’s Rho correlation tests were employed to assess the relationships between damage scores and macrophage subsets.

## Results

### Single-cell RNA-seq and identification of main myeloid cell clusters

Single-cell RNA sequencing (scRNA-seq) was performed of never-frozen kidney immune cells from five AGN patients with active disease, one disease-control with active lupus nephritis (LN), and one nephrectomy control (NC) (**Fig. 1a**). Three patients were PR3-ANCA positive and two patients were MPO-ANCA positive. Due to the necessity of immediate processing, selective sampling was precluded. Nevertheless, all AGN patients had high disease activity scores^21^, histological signs of acute glomerular inflammation. All patients were treated with immunosuppressive medication 2-3 days prior to the kidney biopsy (**Table S1-2**). The LN kidney biopsy was taken pre-treatment and unexpectedly showed a mild proliferative mesangial LN (Class II^19^) with a low activity and chronicity score.

**Fig. 1:**
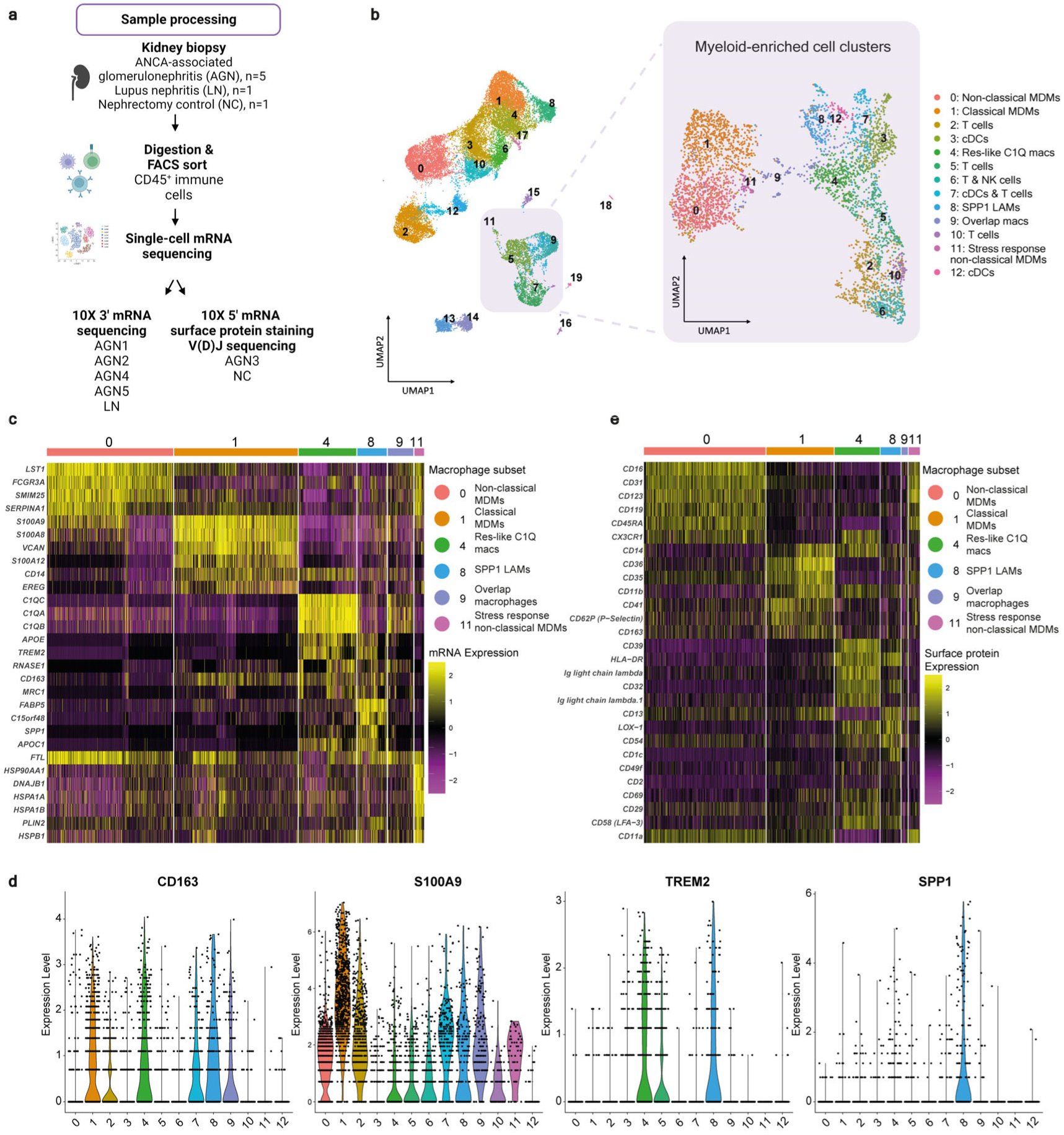
Identification of kidney macrophage subtypes reveal six macrophage clusters. **a**, Flowchart of sample processing. **b**, Uniform manifold approximation and projection (UMAP) visualization of 25,485 single kidney immune cells acquired as in **a**. 20 clusters were identified and annotated using the mature immune kidney cell atlas of Stewart et *al.* 2019 as reference. Myeloid-enriched clusters were selected (located within purple square) and subsequently re-clustered in 13 subclusters containing six macrophage subtypes, which were annotated using canonical cell type markers. **c**, Heatmap showing scaled single-cell mRNA expression of top five markers and a selection of important marker genes per macrophage subcluster. **d**, Violin plots showing high *CD163* expression in macrophage clusters 1,4,8,9, high *S100A9* in cluster 1, and selective *TREM2* and *SPP1* expression in cluster 4 and 8. **e**, Heatmap showing single-cell surface protein abundance per macrophage cluster, validating cluster annotation. MDMs, monocyte-derived macrophages; Res-like macs, resident-like macrophages; LAMs, lipid-associated macrophages; cDCs, classical dendritic cells.

After applying filtering and pre-processing steps, data from a total of 25,485 high-quality immune cells were integrated and clustered for further analysis. Next, we annotated the integrated dataset by mapping it onto the mature immune kidney cell atlas described by Stewart *et al*^15^. Four clusters enriched for myeloid cells were selected and sub-clustered at a low and high resolution distinguishing macrophage clusters and enabling to zoom in on macrophage subtypes, respectively **(Fig. 1b**, **Data S1**). Sub-clustering on low and high resolution resulted in the division of 4,194 cells across five clusters (**Fig. S3, Data S2**) and, correspondingly, 13 clusters (**Fig. 1b**).

### Identification of macrophage subtypes reveals a *SPP1* lipid-associated macrophage cluster

In order to identify (disease-specific) macrophage subtypes, we manually annotated the 13 clusters obtained at high resolution and distinguished six macrophage subtypes; non-classical and classical MDMs (Cluster C0-1), resident-like *C1Q* macrophages (res-like *C1Q* macs) (C4), an *SPP1/TREM2* lipid-associated macrophage (*SPP1* LAM) cluster (C8), a macrophage overlap cluster (C9), and stress response non-classical MDMs (C11) **(Fig. 1b**, annotation and marker expression see **Data S3)**. C0 expressed non-classical monocyte markers^22^ (**Fig. 1c)**, while classical monocyte markers^22^ *CD14*, *CCR2*, *S100A8*, *S100A9*, *S100A12*, *VCAN* and *IL1B* were expressed in C1. C4, res-like *C1Q* macs, abundantly expressed tissue resident markers^15,23^, MHC class II molecules, and complement factors associated with opsonization of apoptotic cells (*C1Q*, *C3*)^24^. Interestingly, cluster 8, *SPP1* LAMs, highly expressed genes associated with lipid scavenging and metabolism^25^, and revealed a transcriptional signature overlapping with gene signatures described in tumor-associated macrophages^26^, profibrotic liver^27^, pulmonary^27^ and kidney macrophages^28^, expressing *SPP1*, *TREM2, LGALS3*, *CD63*, *FABP5*, *LPL*, *CD36*, *CD9*, *APOE*, *APOC1*, *GPNMB*, *PLIN2* and *FN1* (**Fig. 1c-d**). Unexpectedly, the anti-inflammatory macrophage marker *CD163* was not only more highly expressed in *SPP1* LAMs and res-like *C1Q* macs, but also in classical MDMs, often associated with proinflammatory functions (**Fig. 1d)**. The C9 macrophage cluster displayed a mix of cells. C11, primarily originating from the NC sample (**Table S6**), likely represents ischemia-affected non-classical MDMs, expressing stress response markers^30^. Gene-expression based cluster annotation was validated using an antibody cocktail targeting immune cell surface proteins (**Fig. 1e**, **Fig. S4, Data S3**). Interestingly, both res-like *C1Q* macs and *SPP1* LAMs highly expressed surface MHC II proteins (HLA-DR) and LOX-1, a receptor involved in endocytosis of oxidized LDL^31^, highlighting their antigen presenting functions and role in lipid uptake and metabolism. Further analyses were focused on the main four macrophage subsets with >100 cells.

### Enrichment of proinflammatory genesets in classical MDMs and profibrotic genesets in *SPP1* LAMs

Next, we performed geneset enrichment analysis (GSEA), revealing distinctive genesets characterizing each cluster (**Fig. 2a**, **Fig. S5, Data S4**). Non-classical MDMs displayed reduced chemokine (receptor) expression compared to the other three clusters, with upregulation focused on interferon response genesets. Classical MDMs showed significantly higher expression of genesets associated with inflammatory responses, such as TNFα signaling via NFκβ (example gene, log_2_fold change (FC); *TNF*, 1.20; *NFκβ2*; 1.08), degranulation, ROS production, interferon and interleukin-6 and 10, and toll-like receptor (TLR) signaling. Res-like *C1Q* macs exhibited activation of the classical complement pathway, translation, amino acid metabolism and T cells. Chemokine and cytokine receptor interaction genesets were more highly expressed in both res-like *C1Q* macs and *SPP1* LAMs. Interestingly, *SPP1* LAMs showed distinctive expression of genesets related to extracellular matrix (ECM) formation (e.g. epithelial-mesenchymal transition, collagen formation).For example, tissue remodeling-related genes, such as *SPP1* (log_2_(FC) 6.79), *FN1* (5.09), *MMP2* (2.39), *MMP9* (2.21), *LGALS3* (1.93)^33,34^, were upregulated. Also, genesets related to cholesterol homeostasis and uptake were higher expressed in *SPP1* LAMs. Next, we scored cells based on a curated list of genes that regulate ECM production in kidney tissue^35^, which was highest in *SPP1* LAMs (**Fig. 2b**-**c)**. Summarizing, classical MDMs showed specific enrichment of inflammatory pathways, while *SPP1* LAMs exhibited pronounced upregulation of fibrotic genesets.

**Fig. 2:**
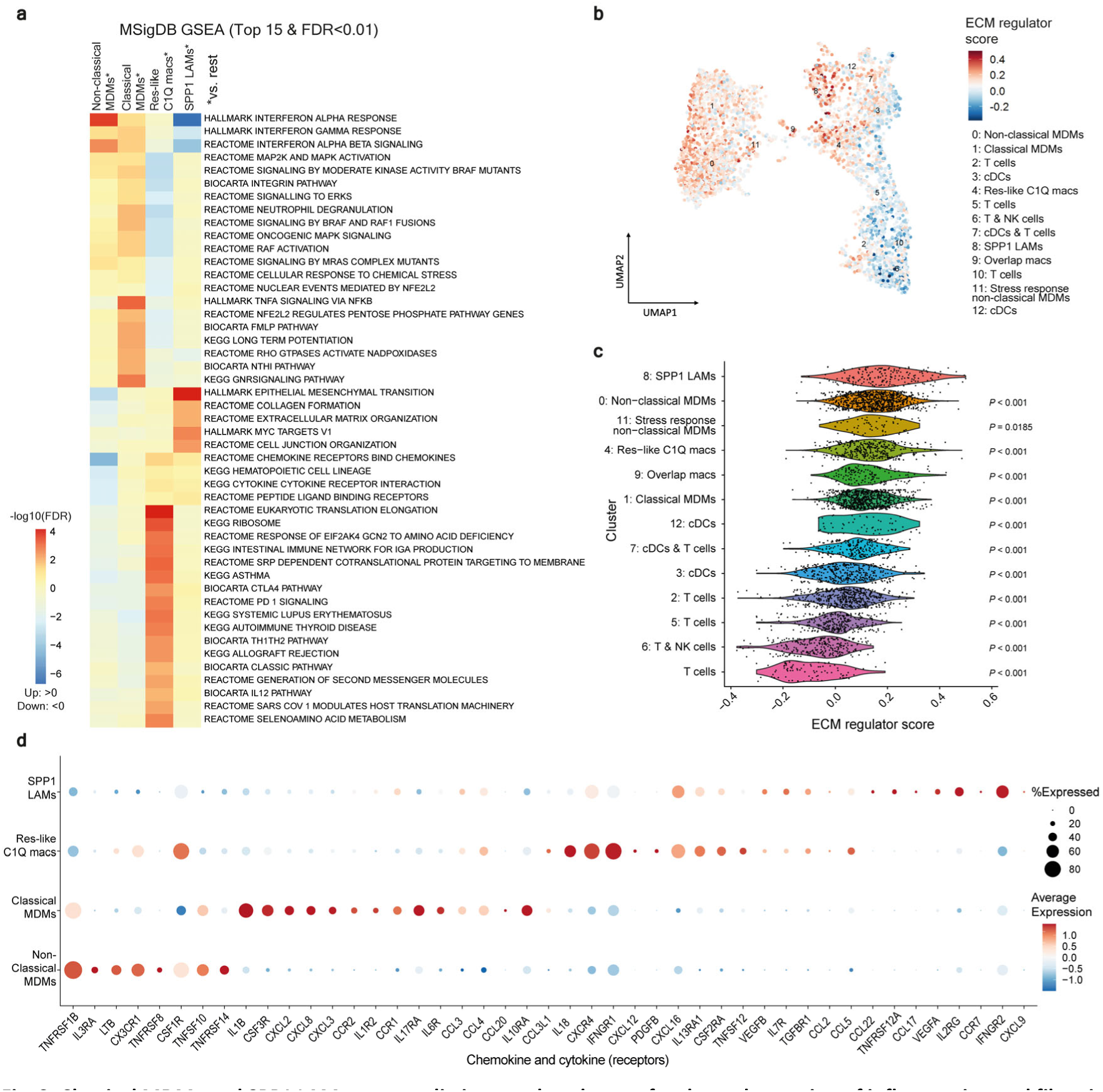
Classical MDMs and SPP1 LAMs express distinct markers known for the orchestration of inflammation and fibrosis. **a**, Geneset enrichment analysis (GSEA) of selected MSigDB geneset collections (Hallmark, BioCarta, KEGG, Reactome) comparing each macrophage subset with the remaining subsets. Heatmap of top 15 significantly (FDR < 0.01) upregulated (red) and downregulated (blue) genesets. **b**, Feature plot of extracellular matrix (ECM) regulator score. **c,** Violin plot showing increased ECM regulator score in *SPP1* LAMs compared to all other clusters. Statistics were calculated using ANOVA with Dunnett’s multiple comparisons using cluster 8 (*SPP1* LAMs) as a reference. **d,** Dot plot illustrating expression of top 15 chemokine (receptor) markers with log2fold change >0.2 per macrophage cluster. MDMs, monocyte-derived macrophages; Res-like macs, resident-like macrophages; LAMs, lipid-associated macrophages; cDCs, classical dendritic cells.

### Classical MDMs express pro-inflammatory cytokines and chemokines involved in the recruitment of neutrophils

Since chemokine and cytokine receptor interaction genesets were more highly expressed in res-like *C1Q* macrophages and *SPP1* LAMs, we compared individual gene expression between the main macrophage subsets (**Fig. 2d**, **Fig. S6).** Compared to the other macrophage subsets, classical MDMs expressed significantly (FDR<0.1) higher mRNA levels of chemokine receptors *CCR1* and *CCR2,* inflammation markers (gene, log_2_(FC); *IL1β*, 1.93; *TNF*, 1.20), and specific mediators for the recruitment of neutrophils (*CXCL2*, 1.77; *CXCL3*, 1.85; *CXCL8,* 2.20)^36^ (**Data S4**). Res-like *C1Q* macrophages highly expressed *CXCR4* and *CSF1R,* the receptor for *CSF1*, an important growth factor for macrophage survival and differentiation^37^. Interestingly, res-like *C1Q* macs exhibited a significantly higher expression of the proinflammatory cytokine *IL18* (log_2_(FC) 1.74). Of note, both *SPP1* LAMs and res-like macrophages significantly expressed *CXCL16* which can act as a receptor for oxLDL uptake^38^. Furthermore, *SPP1* LAMs highly expressed *IFNGR2* (0.55), and exhibited unique coexpression of proangiogenic factors *VEGFA* (2.40) and *VEGFB* (2.05).

### AGN is associated with abundant infiltration of proinflammatory classical MDMs, accompanied by a modest increase in res-like *C1Q* macs and *SPP1* LAMs

We next investigated the expansion of specific macrophage subsets. AGN was associated with an increase in classical MDMs, res-like *C1Q* macs and *SPP1* LAMs, whereas the proportion of non-classical MDMs was reduced compared to the NC sample (**Fig. 3a**-**b**). However, cell type proportions were highly variable and only the proportions of non-classical MDMs were significantly different (FDR=0.004) when taking sample-to-sample variability into account. Next, disease-specific changes within macrophage subsets were investigated. GSEA revealed overall, particularly prominent in MDMs, lower expression of genesets associated with protein metabolism and cellular respiration, and higher expression of genesets related to heme and lipid signalling compared to NC (**Fig. 3c**, **Data S4)**. Given that the LN sample displayed comparable alterations (**Fig. S7a-b)**, these phenomena are likely not exclusive to AGN but might be connected to inflammation in general. Interestingly, both MDM subsets of AGN showed increased chemokine, IL10 and TNFα signaling compared to the NC, in addition to higher TLR, IL1R signaling in classical MDMs, highlighting the inflammatory as regulatory mechanisms of inflammation expressed by their environment. Infiltrating MDMs of AGN (and LN) patients may display an even more pronounced proinflammatory state, since the expression of individual chemokines related to attraction of neutrophils (*CXCL2, CXCL3, CXCL8)* and proinflammatory mediators such as *IL1β* were significantly increased in both AGN MDMs subsets compared to NC. As expected, LN disease control showed increased interferon responses in all macrophage subsets compared to AGN (**Fig. S7b**), but also specific activation of the res-like *C1Q* macs cluster, possibly explained by detection of deposited immune complexes in the kidney tissue. Taken together, AGN is characterized by a markedly elevated proportion of classical MDMs, accompanied by a comparatively smaller elevation in res-like *C1Q* macs and *SPP1* LAMs. MDM gene expression profiles indicate a heightened proinflammatory phenotype in AGN.

**Fig. 3:**
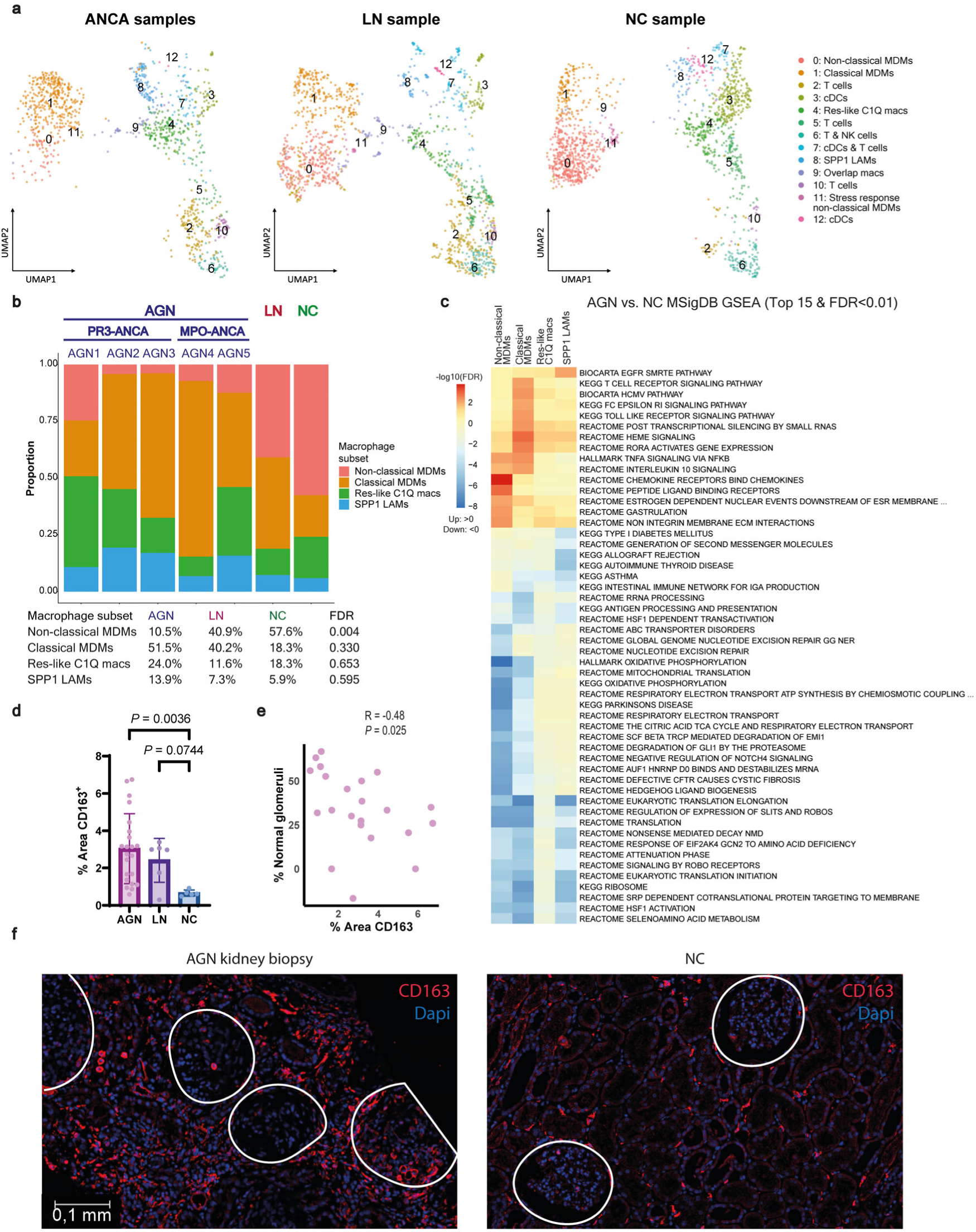
AGN is associated with CD163^+^ macrophage infiltration, consisting of classical MDMs, res-like C1Q macs and SPP1 LAMs. **a,** UMAP visualization of myeloid-enriched cell clusters per disease group. Cluster labels are positioned at the centroid of each cluster. **b**, Stacked-bar plot of macrophage cluster proportions per sample. The table (lower panel) summarizes mean percentage of macrophage clusters per group. Statistics for differences in cell type proportions were calculated using *propeller* with a moderated t-test. The false-discovery rate (FDR) is depicted to account for multiple testing. **c,** Geneset enrichment analysis (GSEA) comparing disease-specific changes within macrophage subsets. Heatmap of top 15 significantly (FDR < 0.01) upregulated (red) and downregulated (blue) genesets. **d-f,** Multi-color immunofluorescence stainings illustrating CD163^+^ macrophages. **(d)** Semi-quantification of the percentage of area positively stained for CD163. Quantitative data are presented as mean ± SD. Statistics were calculated using a Kruskal-Wallis test with Dunn’s multiple comparisons. (**e**) Scatter plot of the Spearman correlation between the percentage of normal glomeruli and percentage area positively stained for CD163 macrophages. (**f**) Representative images showing enhanced infiltration of CD163^+^ macrophages in AGN. AGN, ANCA-associated glomerulonephritis; LN, lupus nephritis; NC, nephrectomy control; PR3, proteinase-3; MPO, myeloperoxidase; MDMs, monocyte-derived macrophages; Res-like macs, resident-like macrophages; LAMs, lipid-associated macrophages.

### Multi-color imaging of kidney biopsies confirms classical MDM infiltration in AGN, in line with histological markers of acute inflammation

To validate differences in kidney macrophage composition and explore histological correlations, multi-color immunofluorescence stainings of kidney tissue were performed in an extended cohort (**Table S7).** Our single-cell analysis revealed high *CD163* mRNA expression across classical MDMs, res-like *C1Q* macs and *SPP1* LAMs (**Fig. 1d)**. As anticipated from these data and existing literature^5,39^, CD163^+^ macrophages were significantly increased in AGN kidney biopsies (*P*=0.0036) (**Fig. 3d)** and correlated negatively with normal glomeruli (**Fig. 3e**-**f**). CD163^+^ macrophages infiltrated the interstitium, encircling inflamed glomeruli, whereas in NC biopsies, CD163^+^ macrophages were uniformly dispersed (**Fig. 3f)**. Macrophage markers were chosen based on scRNA-seq data (**Fig 1c**-**d)** (Classical MDMs: *S100A9*, res-like *C1Q* macs : *TREM2*, *SPP1* LAMs*: TREM2, PLIN2* and *SPP1)*. S100A9/CD163 classical MDMs were significantly increased in AGN (*P*= 0.0384) (**Fig 4a)**, notably higher in samples with top activity scores (**Fig. 3b**), and were abundantly present within patchy infiltrates in the interstitium and cellular crescents (**Fig. 4b,e**, **Fig. S8)**. Flow cytometric analysis of PBMCs showed a similar increase in the classical monocyte subset (CD14^++^ CD16^-^) during active disease, while non-classical monocytes (CD14^+^ CD16^++^) were significantly decreased (**Fig. 4c**).

**Fig. 4.**
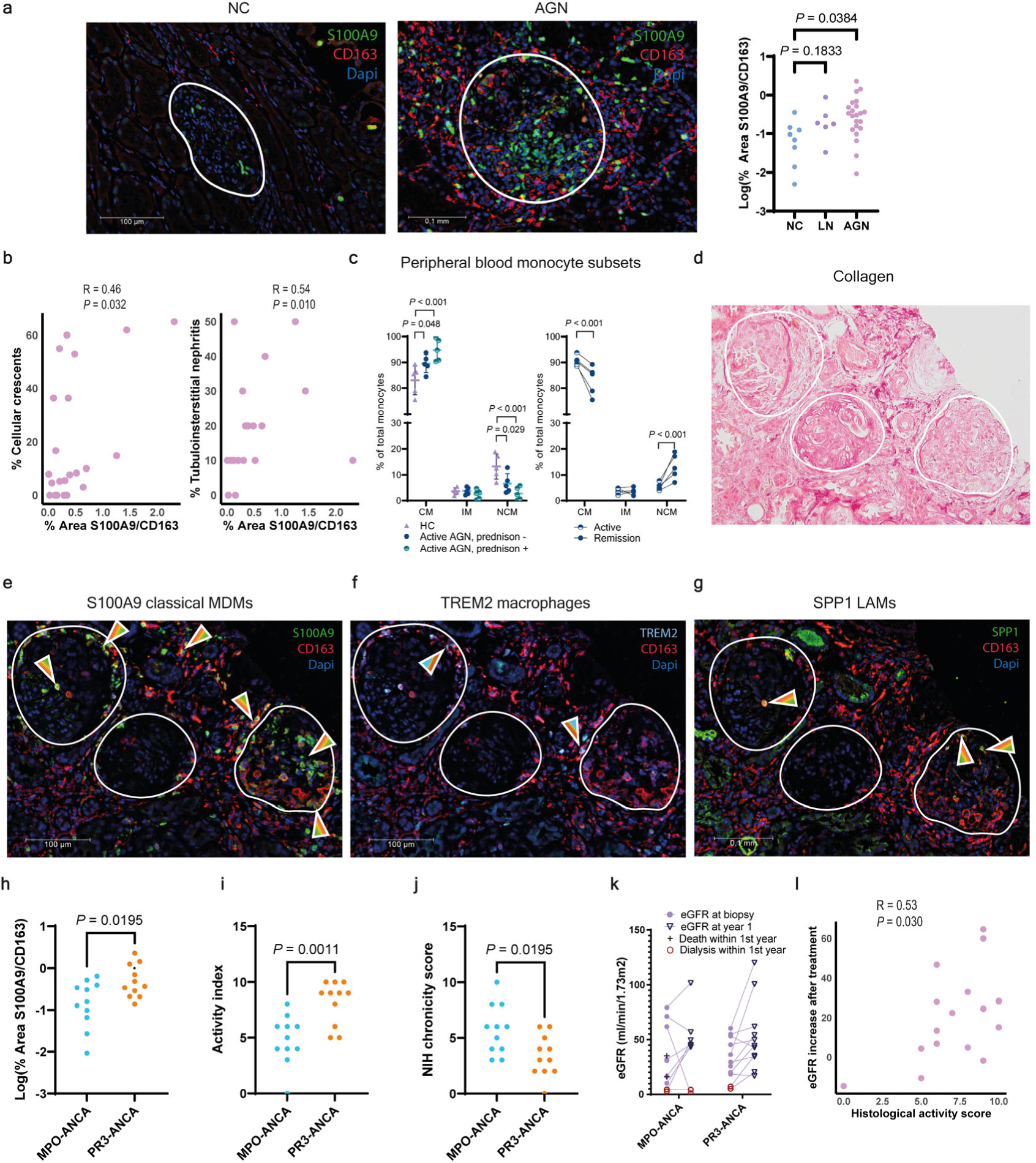
Enhanced infiltration of classical MDMs correlates with histological signs of inflammation and the PR3-ANCA serotype. **a**, Multi-color immunofluorescence stainings of CD163/S100A9 macrophages (representing classical MDMs) in kidney biopsies of AGN, LN and NC. Semi-quantification shows significantly increased CD163/S100A9 macrophages in AGN. Quantitative data were log-tranformed and individual data are presented. Statistics were calculated using a Brown-Forsythe ANOVA with Dunnett’s T3 multiple comparisons. **b**, Scatter plot of the Spearman correlation between the percentage of cellular crescents (left) and percentage of tubulointerstitial nephritis (right) and percentage area positively stained for S100A9/CD163 macrophages. **c**, Flowcytometric analysis of peripheral blood monocyte subsets. Cross-sectional comparison of cell type proportions (left) between patients with active AGN with and without concurrent prednisone treatment compared to matched healthy controls. Paired analysis of cell type proportions (right) during active and stable disease. Monocyte subsets are expressed as a percentage of total monocytes. Quantitative data are presented as mean ± SD. Statistics for differences in cell type proportions were calculated using *propeller* with a moderated t-test (left panel, unpaired; right panel, paired), Benjamini-Hochberg FDR corrected. **d-g,** Picro Sirius red and multi-color immunofluorescence stainings of CD163, S100A9, TREM2 and SPP1, in human kidney biopsies of AGN patients. Glomeruli are delineated by white circles. (**d**) Picro Sirius Red staining highlighting presence of fibrosis (dark red). (**e**), shows S100A9/CD163 macrophages located in the interstitium and encircling active AGN lesions. (**f)** TREM2/CD163 and (**g**) SPP1/CD163 macrophages are located in areas of inflammation. **h-l,** comparisons between AGN serotypes (**h**) Semi-quantification of the percentage area positively stained for S100A9/CD163 macrophages (representing classical MDMs) per serological subset. Quantitative data were logtransformed and individual data are presented. (**i**) Histological activity score is higher in PR3-ANCA (n=11) compared to MPO-ANCA (n=11), while (**j**) the chronicity score is lower. (**h-j**) Statistics were calculated using an unpaired t test (**h-i**) and Mann-Whitney test (**j**). **k,** Longitudinal follow-up of estimated glomerular filtration rate (eGFR) per ANCA subtype. **l,** Scatter plot of the Spearman correlation between the delta eGFR within the first year and histological activity index. CM, classical monocyte; IM, intermediate monocytes; NCM, Non-classical monocytes. MDMs, monocyte-derived macrophages; LAMs, lipid-associated macrophages. MPO, myeloperoxidase; PR3, proteinase 3.

SPP1/TREM2/CD163 macrophages (*SPP1* LAMs) and TREM2/CD163 (res-like *C1Q* macs) were observed both within and around the inflamed glomeruli (**Fig. 4e**-**g, Fig. S8-9**), although numerically much less than the S100A9 macrophages. Picrosirius Red staining suggested contingency between the stage of fibrosis and macrophage infiltration. In end-stage sclerotic glomeruli, macrophages were absent, while in and around glomeruli displaying concurrent inflammation and/or early fibrotic changes, both S100A9/CD163 and SPP1/TREM2/CD163 macrophages were present (**Fig. 4d-g**, **Fig. 5)**, suggesting a link between inflammation-induced epithelial injury and early fibrotic changes^40^. Finally, macrophage composition and phenotype were assessed between ANCA subtypes. Comparing MDM phenotypes showed increased interferon signaling in MDMs of the two MPO-ANCA patients (**data S4)**. In the extended histological cohort, the percentage of classical S100A9/CD163 MDMs, along with the activity index score, were significantly increased in kidney biopsies of PR3-ANCA patients (**Fig. 4h**-**i**), while the histological chronicity score was significantly lower compared to MPO-ANCA patients (**Fig. 4j)**. These findings confirm the link of classical MDMs with acute inflammation, and therefore higher prevalence in the PR3-ANCA subtype^3^. Since persistence of dialysis and death were only present in the MPO-ANCA subgroup (**Fig. 4k)**, and the histological activity index score was positively correlated with recovery of kidney function within the first year of treatment (**Fig. 4l)**, this may highlight the capacity to effectively target classical MDMs prior to the occurrence of irreversible chronic damage in PR3-AGN.

**Fig. 5:**
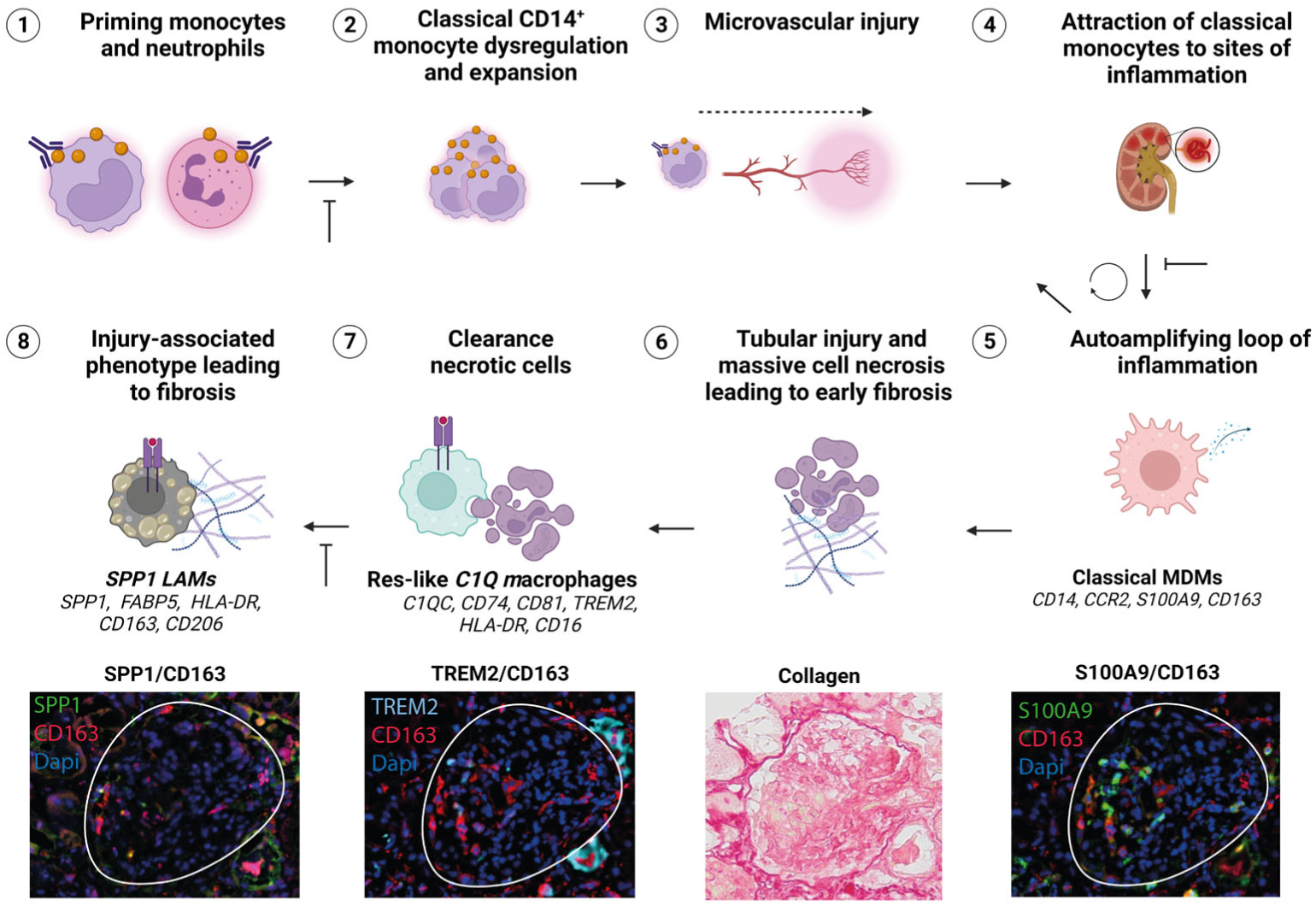
Kidney macrophages might enhance inflammation and fibrosis in AGN. Illustration shows proposed chain of events in development of vasculitic lesions and long-term fibrosis based on present findings and literature. (1-3) ANCAs are generated and bind primed peripheral blood monocytes and neutrophils causing classical monocyte dysregulation, expansion and microvascular inflammation in kidney glomeruli. (4) Classical monocytes migrate into the kidneys, and (5) differentiate locally into classical MDMs where they amplify inflammatory cascades by production of inflammatory chemokines and neutrophil-recruiting cytokines. (6) Resulting tubular injury induces fibrotic cascades early on and results in massive cell necrosis, (7) which attracts resident *TREM2* macrophages to clear necrotic debris. (8) Ultimately, these cells are overwhelmed and start to contribute to fibrosis and express an injury-associated phenotype including *SPP1*. These findings shed new light on the complexity of the macrophage immune response in AGN and indicate a complementary role for macrophages in neutrophil-mediated disease initiation and progression.

## Discussion

Kidney tissue macrophage function in AGN has been postulated to encompass both the promotion and resolution of inflammation. Here, we define a spectrum of kidney macrophages, ranging from monocyte-derived macrophages (MDMs), resident-like macrophages (res-like *C1Q* macs) and an *SPP1* lipid-associated macrophage subtype (*SPP1* LAMS). We illustrate that classical MDMs, res-like *C1Q* macs and *SPP1* LAMs are enriched in kidneys of AGN patients, and express remarkably divergent immune response phenotypes. While classical MDMs highly expressed genes related to inflammation, the expression profile of *SPP1* LAMs was related to extracellular matrix organization and fibrosis. These findings shed new light on the sophisticated immune response in AGN.

Notably, AGN kidneys were enriched for profibrotic *SPP1* kidney macrophages expressing a transcriptional signature associated with lipid scavenging, lipid metabolism and tissue-remodeling, such as *TREM2, MMP7*, *MMP9*, *CTSD*, *TIMP1*. Osteopontin (SPP1) is a glycoprotein widely expressed in the kidney by tubular and immune cells, and mediates multiple physiological and inflammatory processes^41,42^. Osteopontin has been identified as a biomarker for disease activity in auto-immune kidney diseases^43,44^, and is linked to kidney failure^45^ and fibrosis^42,46^. Our results support previous reports identifying *SPP1/TREM2* macrophages as cross-tissue regulators of fibrosis in various tissues by the induction of an injury associated phenotype^28,47^. Macrophages engaged in homeostatic processes have been shown to transit towards a fibrotic polarization state, likely in situations with unresolving inflammation. Indeed, in AGN, *SPP1* was linked with collagen and fibronectin expression, and glomerular damage indicated by breach of BC^48^, facilitating the entry of CD8^+^ T cells into the glomerulus^49^. Eventually, *SPP1* macrophages can contribute to fibrosis by ECM formation or fibroblast interaction via FN1, SPP1 and SEMA3^28,51^, which were all upregulated. Taken together, profibrotic *SPP1* LAMs seem to emerge within a highly inflammatory environment, where macrophages attempt to remove necrotic cells, because of unresolved inflammation eventually enforcing a fibrotic cascade.

Next, AGN kidneys demonstrated an increased resident-like *C1Q* macrophage population, associated with both an immunosuppressive and inflammatory phenotype. In homeostasis, C1Q^+^ macrophages serve an important immunoregulatory function by phagocytosis of C1Q opsonized apoptotic cells and reduction of pro-inflammatory mediators^24,52,53^. TREM2 and SPP1 expressing macrophages were mainly located intra- and periglomerularly, suggesting their attraction by glomerular inflammation to clear necrotic debris. This is in line with earlier spatial transciptomic findings showing increased *SPP1* and *C1Q* expression in glomeruli of ANCA patients with rupture of BC^48^. Res-like *C1Q* macs highly expressed *CSF1R, IL18* and *HLA-DR*, indicating enhanced proliferation through autocrine signaling, but also inflammatory responses. Indeed, in ANCA-associated vasculitis, IL-18 was upregulated in interstitial kidney macrophages, and primed neutrophil responsiveness^54^.

Most importantly, infiltrating classical MDMs highly expressed markers of inflammation and immune cell recruitment in kidneys of AGN patients, whereas non-classical MDMs were present in larger numbers during homeostasis. These results confirm findings of a murine study transplanting hematopoietic cells in MPO immunized mice, highlighting the necessity of CCR2^+^ classical monocytes for AGN development^55^ . Using specific knock-out models, CSF2 was shown to stimulate monocytic IL1β release and Th17 polarization by CM. The classical MDM subset indeed exhibited high expression of CCR2 and IL1β, but not IL-6, known for T_h_17 polarization^56^. Besides these proinflammatory cytokines, classical MDMs expressed S100 molecules, and chemokines known for neutrophil recruitment. Research has demonstrated that neutrophils play a crucial role in the pathophysiology of ANCA-induced endothelial damage^57^. Release of these mediators may enhance the downward spiral of escalating neutrophil-mediated inflammation^57^, although functional tests have not been performed. The correlation between classical MDM infiltration and disease activity was also shown by elevated serum calprotectin levels, a heterodimeric complex of S100A8 and S100A9, during active disease and calprotectin staining in active glomerular lesions^58^. Spatial transcriptomics indeed confirmed the relation between macrophages and acute inflammation by the expression of CD163 in early glomerular lesions^48,59^. Utilizing a multi-color microscope enabled us to precisely co-localize and differentiatiate S100A9 expression on macrophages from other immune cells, such as neutrophils.

Observing an analogous trend in the expansion of peripheral blood classical monocytes during active disease, classical monocyte dysregulation seems to be an important event in the pathogenesis of AGN. This was also highlighted by a recent sc-RNAseq paper^60^, confirming the increase of classical CD14^+^ monocyte subsets, including an interferon signature (ISG) expressing subset and an immature activated classical monocyte subset. The activated subset closely resembled the expression profile of classical MDMs, including top markers *S100A9*, *S100A8*, *VCAN* and *CTSD*^60^, and was clinically related to relapses, whereas the ISG subset was related to kidney disease. While classical CD14^+^ monocyte infiltration is related to disease activity in both serological subsets, it could be speculated that varying prevalences of the ISG and activated classical monocyte phenotypes could potentially give rise to phenotypical differences. PR3-ANCA generally have a higher risk of relapses^3^ and had higher S100A9^+^ classical macrophage infiltration, whereas kidney involvement is more frequent in MPO-ANCA vasculitis^3^ and classical MDMs of MPO-AGN patients showed higher interferon responses compared to PR3-AGN. Since peripheral blood monocytes of MPO-AAV patients show a similar upregulation of interferon responses (data submitted^61^), observations could represent actual pathological distinctions specific to different serotypes, rather than variations related to the stages of the disease. However, the limited sample size and heterogeneity of the samples included in our study precluded a more in-depth investigation of these potential causes, further research investigating serotype-specific effects are needed.

The use of scRNA-seq enabled us to profile kidney immune cells in AAV, uncovering pathways and markers involved in the local immune response. While the use of freshly retrieved unfrozen cells avoided bias due to preservation methods, this method inherently limited sample size and sample selection. Therefore, substantial heterogeneity between samples was noted, especially variation in disease severity, chronicity, and use of immunosuppressive treatments. Validation studies used specific markers, but lacked functional assays, limiting causal conclusions. Despite these limitations, the findings provide valuable insights into the kidney immune response, and provides a foundation for future research to expand upon.

This deeper understanding of macrophage dynamics may offer new avenues for developing precision treatments for AGN and autoimmune kidney diseases in general. Research should focus on targeting classical monocytes and classical MDMs in early phases to dampen acute inflammation before the start of fibrotic cascades (**Fig. 5)**. Concurrently, targeting *SPP1* macrophages may reduce long-term fibrotic processes, possibly also in other non-immune mediated kidney diseases. Current glucocorticoid regimens are likely very effectively targeting classical MDMs^62^, whereas a more precise and promising avenue could involve a CD163-targeted dexamethasone delivery system^63^.

Ultimately, specific targets to modulate the inflammatory cascade and resulting kidney fibrosis could highly benefit kidney disease in general.

In summary, our study highlights the expression of genes associated with inflammation and fibrosis by kidney macrophages in AGN. These findings shed new light on the complexity of the macrophage immune response and suggest a complementary role for macrophages in neutrophil-mediated disease initiation and progression. A deeper understanding of macrophage dynamics could offer new avenues for developing precision treatments aimed at mitigating the onset and impact of autoimmune kidney diseases ultimately paving the way for more effective management strategies in its early phases.

### Disclosures

JK has received consultancy fees from Aiosyn BV, Alentis Pharmaceuticals AG, Center for Human Drug Research Leiden and is a Central Biopsy Review coordinator for Novartis AG (SIRIUS-LN randomized trial). FJB and LV have acted as member of the scientific advisory board of the Dutch Kidney Foundation.

## Acknowledgements

We would like to thank the MOMA study collaborators: N. van der Bom-Baylon, T. Dekker, dr. E.L Penne, B. Hooibrink. Fig. 1a, Fig. S2a, and Fig. 5 were created with BioRender.com. Date were presented at the ASN Kidney Week 2023.

## Ethics statement

The research protocol was approved by the local ethics boards in accordance with the Declaration of Helsinki, and subjects gave their written informed consent.

