## Supplementary material for "Infiltrative classical monocyte-derived and *SPP1* lipid-associated macrophages mediate inflammation and fibrosis in ANCA-associated glomerulonephritis"

**Supplemental document**

### Supplementary Methods

#### Study subjects and kidney tissue acquisition

Renal tissue was acquired at two hospitals in the Netherlands. According to the Declaration of Helsinki, the research protocol was approved by the locally appointed ethics committees and written informed consent was obtained from the subjects. Kidney tissue from AGN patients (n=5) and a lupus nephritis (LN) patient (n=1) were aquired during a biopsy procedure for a clinical indication. Needle-core biopsies were obtained for histopathological examination, and an additional pass was performed to retrieve kidney tissue for scRNA-seq with a maximum total passes of 3 to ensure the safety of participants. Healthy kidney tissue (n=1) was obtained from a kidney that was surgically removed due to a (non-invasive) papillary urothelial carcinoma. Since we expected lower immune cell infiltration in the healty kidney compared to the AGN samples, a total amount of 50 passes were obtained from the healthy cortex to ensure sufficient immune cell recovery. The samples are referred to as AGN, LN and NC, respectively. All AGN patients had high disease activity scores (median activity score of 16 assessed by the Birmingham Vasculitis Activity Score (BVASv3)[14], histological signs of acute glomerular inflammation (median cellular crescent proportion of 50%), and were treated with immunosuppressive medication 2-3 days prior to the kidney biopsy (80% received methylprednisolone therapy). The LN kidney biopsy was taken prior to start of treatment and histological examination revealed a mild proliferative mesangial LN, defined as a class II lupus nephritis according to the International Society of Nephrology/Renal Pathology Society classification system [15] with a low activity (1/24) and chronicity score (3/12) (**Table S1-2**).

#### Kidney tissue processing

Immediately after collection, kidney tissue was processed into a single-cell suspension using a modified protocol described earlier [16]. Briefly, kidney tissue was placed into a 20 ml preservation solution on ice (Iscove's Modified Dulbecco's Medium (IMDM, Thermo Fisher Scientific, Gibco, 12440061) supplied with 10% Fetal Calf Serum (FCS, VWR International BV) + 0,38% TNC (Merck, 1.06448.0500) + 0.00036(v/v)% β-mercaptoethanol (Merck, M-7154)). Kidney specimens were washed twice with cold Phosphate-buffered saline (PBS) and cut into 6-7 pieces. Kidney pieces were transferred into 2ml of preheated (37 °C) digestion medium (50 KU/mL DNAse I type IV + 0.5 mg/mL collagenase type IV + 60 mg/mL BSA + 20 µL/mL FCS + 0.025 M TRIS + penicillin streptomycin in HBSS), and incubated for 20 min at 37 °C while shaking. The digestion process was stopped by adding cold preservation solution and the resulting cell suspension was passed through a 70 um cell strainer (Falcon, 734-0003) to remove debris and acquire a single-cell suspension. Since a large amount of kidney tissue was processed in the healthy kidney sample to ensure sufficient immune cell recovery, standard density gradient centrifugation was used to decrease sorting time according to manufacturer's protocol (Lymphoprep, Abbott Diagnostics Technologies AS, AXI-1114547).

**Flow cytometric cell sorting**

The kidney single-cell suspension was sorted using a 4-color flow cytometry panel to isolate living, $CD45^+$ immune cells (**Supplementary Fig. S1**). Antibodies for cell sorting include anti-CD45 and a viability dye, while CD3 and CD14 antibodies were used to estimate T cell and monocyte/macrophage proportions, respectively, at time of cell sorting (**Table S3-4**). Kidney cells were incubated with these surface antibodies for 30 min at 4 °C in the dark, washed once with RPMI 1640 medium (Thermo Fisher Scientific, Gibco, 11875093) supplied with 5% FCS and filtered (Filcon 50 um syringe, BD, 340601) to remove debris. Single cells were sorted on a BD FACSAria IIu SORP 3- or 4- laser sorter and $eFluor506UV^-$ $CD45^+$ living immune cells were taken up in cold 1mL RMPI/10% FCS medium. Flow cytometric quantification of cell populations was performed using FlowJo 10.0.7.

**Surface marker labeling with TotalSeq™-C Human Universal Cocktail for donors AGN3 and NC**

To aid in cluster annotation and multi-omic characterization of kidney immune cells, surface protein abundance and T and B cell receptor repertoires were determined in two samples (AGN3, NC). These samples were incubated with an oligo-antibody TotalSeq-C cocktail containing 130 unique cell surface antigens after cell sorting. The cocktail (Biolegend, 399905) was reconstituted according to the manufacturer's protocol. Kidney cells were first incubated with Trustain FcX (Biolegend, 422301) shaking for 5 min at 4 °C in the dark to reduce background labeling and then incubated with the antibody cocktail shaking for 15 minutes at 4 °C in the dark.

**Loading of cells and single-cell library preparation**

Immediately after cell sorting (AGN1, AGN2, AGN4, AGN5 and LN) or after labeling with TotalSeq-C Human universal Cocktail (AGN3 and NC), cells were washed twice and resuspended in PBS + 0.1% BSA at a target concentration of 1000 cells/ul. Next, cells were loaded onto a Chromium chip (10x Genomics) for a target recovery of 8,000 cells per sample. This corresponds to a >99% probability of capturing at least 50 cells from each kidney immune cell type assuming 21 cell types in total and a minimum proportion of the rarest cell type of 0.01 (https://satijalab.org/howmanycells/). Considering loss by inadequate capturing of cells, generation of doublets, and filtering steps, we determined a target loading number of 14,000 cells per sample (https://satijalab.org/costpercell/). TCRαβ and BCR libraries were prepared with the V(D)J enrichment Kit from 10x Genomics (Chromium Single Cell 5' Feature Barcode Library Kit v1.1, 10x Genomics, PN-1000080). Since T and B cell receptor repertoire analysis requires 5 'sequencing, cDNA synthesis, amplification, and sequencing libraries were made using either the 10X Genomics single-cell 3' Reagent (Chromium Next GEM Single Cell 3' Library & Gel Bead Kit v3.1, 10x Genomics, PN-1000128) or the 5' Reagent (Chromium Next GEM Single Cell 5' Library & Gel Bead Kit v1.1, 10x Genomics, PN-1000167), depending on the sample **(Table S4)**. Our initial strategy involved employing 5' sequencing for all kidney immune cells. However, this approach faced challenges due to the loss of three samples when combining 5' sequencing with surface marker labeling. Consequently, we opted for 3' sequencing for subsequent samples. Gene expression libraries were sequenced on an Illumina NovaSeq 6000 platform at a target depth of 30,000 reads per cell. For AGN3 and NC, surface protein libraries and V(D)J libraries were pooled with gene expression libraries at a target depth of 10,000 reads/cell (surface proteins) or 5,000 reads/cell (V(D)J).

**Single-cell data pre-processing and analysis**

Demultiplexing of raw base call (BCL) files and conversion to FASTQ files were performed with the Cell Ranger pipeline (10x Genomics, version 6.1.2) using 'cellranger mkfastq'. Read trimming, alignment to the human reference genome and transcriptome (10x Genomics; GRCh38-2020-A), barcode correction, UMI counting and filtering of empty barcodes were performed using 'cellranger count' for samples with only a gene expression library. For samples with gene expression, surface protein and V(D)J libraries 'cellranger multi' was used to combine these steps with UMI counting for the surface proteins, V(D)J sequence assembly, contig annotation and clonotype counting (using refdata-cellranger-vdj-GRCh38-alts-ensembl-5.0.0 as reference). The expected number of cells was set to 8000.

Resulting filtered feature-barcode matrices were imported in R using the Seurat package (v4.1.1). For each cell barcode the number of expressed genes in the list of 98 housekeeping genes of Tirosh et al. (2016) [17] and the percentage of reads mapping to mitochondrial genes were determined. To remove cell barcodes of low quality or potential doublets, we excluded those with <200 or >=6.000 detected genes, <=55 expressed housekeeping genes or a percentage of mitochondrial reads >=10%. Only genes with at least one feature count in >=3 cells were used in the analysis. Normalization and variance stabilization of the resulting gene-level count data were performed using regularized negative binomial regression, while regressing out the percentage of mitochondrial reads in a second non-regularized linear regression (Seurat; function SCTransform, vars.to.regress = "pct.mito"). Surface protein counts were normalized using a centered log ratio (CLR) transformation across cells (Seurat; function NormalizeData, normalization.method="CLR", margin=2)

Doublet removal was performed in two steps. First, for the two samples with a V(D)J library, barcodes with both high-confidence B and T cell receptors were removed. Second, we ran DoubletFinder for each of the gene expression libraries separately. This involved further processing of each Seurat object by selecting the 2000 most variable genes (Seurat; function FindVariableFeatures) and performing principal component analysis (PCA) (Seurat; function RunPCA, npcs=100). The following non-default values for the DoubletFinder parameters (function DoubletFinder_v3) were used: number of principal components (PCs), nPC=1:35; principal component neighborhood size, pK=0.09; logical indicating that SCTransform was used, sct=TRUE; total number of doublet predictions, nExp=0.075*number of barcodes. Here, 0.075 was chosen since this is the expected doublet rate when loading 14,000 cells. Barcodes classified as doublets by DoubletFinder were removed.

Integration, clustering and annotation were performed in an iterative manner. First, we annotated each individual sample with SingleR using the mature full kidney cell atlas (https://cellgeni.cog.sanger.ac.uk/kidneycellatlas/Mature_Full_v3.h5ad) of Stewart et al. [18] as a reference and removed cells classified as proximal tubule cells. Samples of the individual donors were integrated using Seurat's anchor-based integration [19] with the standard workflow for data normalized with SCTransform (functions SelectIntegrationFeatures, PrepSCTIntegration, FindIntegrationAnchors, IntegrateData). We selected 3000 features for integration (function SelectIntegrationFeatures) and used dimensionality reduction with reciprocal PCA and AGN3 as a reference for finding the integration anchors (function FindIntegrationAnchors, reduction=”rpca”). The top-30 PCs calculated on the integrated data were used as input for non-linear dimensionality reduction using UMAP (Seurat; function RunUMAP) and for graph-based clustering (Seurat; functions FindNeigbours, FindClusters) with resolution 0.6. Subsequently, we annotated the integrated dataset by mapping it onto the mature immune kidney cell atlas (https://cellgeni.cog.sanger.ac.uk/kidneycellatlas/Mature_Immune_v2.1.h5ad) of Stewart et al. [18] as reference. First, the original gene-level counts of the reference data were normalized using SCTransform and the top-30 PCs were calculated. Next, anchors between the reference data and our integrated data were determined (Seurat; function FindTransferAnchors). The set of calculated anchors was then used to transfer cell type annotation from the reference to our integrated dataset (Seurat; function TransferData). Finally, we determined the UMAP model of the reference and projected the integrated dataset onto the reference UMAP (Seurat; function MapQuery). Clusters at resolution 0.6 enriched for myeloid cells, that is containing ≥ 5% of all myeloid cells, were selected (**Table S5)**. The resulting integrated myeloid-enriched dataset contained 4,194 cells and was used as the basis for all analyses below.

We then renormalized the gene and surface protein counts for each sample separately, using the same methods and parameters as above. Samples of the individual donors were again integrated using Seurat’s anchor-based method as above, except that now dimensionality reduction with canonical correlation analysis (CCA) was used and no reference was specified (Seurat; FindIntegrationAnchors, default parameter values). The integrated dataset was clustered as above at resolutions 0.3 and 1. Genes differentially expressed (DE) between a cluster and all other clusters were determined on the SCTransform-normalized data using logistic regression with ‘donor’ as latent variable (Seurat; function FindAllMarkers, test.use=”LR”, min.pct= 0.25) for the clusters at resolution 1. Cell type annotation of the clusters was performed based on the occurrence of canonical cell type markers in the top of the ranked list of cluster markers. Differentially expressed surface proteins were determined on the CLR-transformed data using a Wilcoxon rank sum test (Seurat; function FindAllMarkers, test.use=”wilcox”, min.pct= 0.25). Marker genes and surface proteins were visualized using heatmaps and dot plots (Seurat; functions DoHeatmap, DotPlot).

Genes differentially expressed between different conditions for cells of the same type were determined using a pseudobulk-based analysis [20]. Here, we aggregated the raw counts per combination of donor and cell type to construct the pseudobulk samples. Statistical analyses were performed using the edgeR and limma R/Bioconductor packages. Genes with more than 2 counts-per-million reads (CPM) in 1 or more of the samples were kept and were reannotated using Ensembl (v109) with biomaRt. Count data were transformed to log2-counts per million (logCPM), normalized by applying the trimmed mean of M-values method and precision weighted using voom. No sample outliers were identified based on library size or normalization factors. Differential expression was assessed using an empirical Bayes moderated t-test within limma’s linear model framework, including the precision weights estimated by voom. A consensus intra-donor correlation was estimated for pseudobulk samples from the same donor but from a different cell type (limma; function ‘duplicateCorrelation’) and included in the linear model fit. The Benjamini-Hochberg false discovery rate (FDR) was used for multiple testing correction of the resulting *P*-values. An adjusted *P*-value <0.1 was considered significant.

Geneset enrichment analysis (GSEA) was performed using CAMERA (limma package) with a value of 0.01 for the inter-gene correlation, using selected geneset collections (Hallmark collection and the BIOCARTA, KEGG and Reactome subsets of the C2 collection) from the Molecular Signatures Database (MSigDB; v2023.1.Hs). *P*-values were calculated using a two-sided directional test (direction of change, 'up' or 'down') and corrected for multiple testing using the Benjamini-Hochberg FDR. An adjusted *P*-value <0.1 was considered significant. Results for selected genesets and comparisons were visualized using heatmaps and enrichment networks. For the latter, aPEAR was used to group similar genesets together and to highlight the most important themes in the geneset enrichment results. An FDR < 0.1 was used to select genesets per comparison. Default settings for the calculation of the similarity matrix between genesets ('jaccard'), clustering ('markov'), and naming of the clusters ('pagerank') were applied.

Myeloid-enriched cells were scored (Seurat; function AddModuleScore) based on a curated list of genes that regulate extra-cellular matrix (ECM) production in kidney tissue retrieved from MatrisomeDB (http://matrisomedb.pepchem.org/; selection: ECM Regulators – Human - kidney).

Differences in cell type proportions between conditions were assessed using *propeller* [21]. First, proportions were transformed using a logit function in order to reduce heteroscedasticity. Next, we performed moderated t-tests to test whether the transformed proportions were significantly different between groups. Resulting *P*-values were corrected for multiple testing using the Benjamini-Hochberg FDR. Analyses were performed using R v4.1.0, Bioconductor v3.13 and Seurat v4.1.1.

**Flowcytometric analysis of peripheral blood monocytes**

Blood was collected from 10 patients with ANCA-associated vasculitis during active disease, 5 follow-up samples during stable disease and 5 age-and-sex matched healthy controls. PBMCs were isolated using standard density gradient centrifugation and cryopreserved until analysis. Measurements were performed on a BD FACSCanto flow cytometer. Per sample, 6 panels were measured using 0.25 x $10^6$ PBMCs per panel. The volume of each staining reaction remained unchanged, as did the concentrations of antibodies. PBMCs were incubated with a fixable viability dye eFluoro506 and 6 panels with surface antibodies (**Table S3**) for 30 min at 4°C in the dark, and washed twice before measurements. Gating strategy to distinguish monocytes and monocyte subsets is shown in **Fig. S1**. The proportion of monocyte subsets was evaluated for each panel, and the mean percentage was calculated for each participant. Data were analyzed using FlowJo version 10.0.7 (FlowJo, Ashland, OR, USA) and Graphs were made using Graphpad Prism version 9.3.1.471 (GraphPad Software, La Jolla, CA, USA). Differences in proportions of peripheral blood monocyte subsets between conditions were assessed using *propeller* [21]. First, proportions were transformed using a arcsin square root function in order to reduce heteroscedasticity. Next, we performed moderated t-tests (either paired [active versus remission] or unpaired [rest]) to test whether the transformed proportions were significantly different between groups. Resulting *P*-values were corrected for multiple testing using the Benjamini-Hochberg FDR.

**Multi-color immunofluorescence stainings of kidney tissue extended cohort**

The presence of *SPP1/ TREM2/PLIN2* LAMs and *S100A9* classical MDMs was validated using immunofluorescence stainings of an extended cohort of 36 kidney biopsies (22 AGN, 6 LN, 8 NC). 4-µm-thick paraffin-embedded kidney sections were freshly cut and dried overnight in a 37 °C oven. The paraffin-embedded tissue sections were deparaffinized by sequential incubation in xylene and a graded ethanol series. Subsequently, tissue sections were rehydrated in distilled water (aqua dest). Antigen retrieval was performed by boiling the samples in a 10 mM Tris/EDTA buffer in an autoclave for 10 minutes at 100 °C. Immunostaining was carried out using antibodies depicted in **Table S3**. Primary antibodies were stained overnight at 4°C, followed by a stepwise 60 min incubation of the secondary antibodies to avoid cross-binding of anti-goat AF750 with secondary goat antibodies (Goat AF633, Goat AF568). Kidney biopsies were mounted in Aqua Poly/Mount (Polysciences, Inc.) and visualized. Whole-biopsy multi-color images were taken using the Thunder Microscope system (Leica). The settings, including laser intensity and exposure time, remained consistent throughout the acquisition of all images. Quality of the staining was assessed and 1 biopsy had to be excluded from all analyses due to excessive non-specific staining. To visualize the relationship between macrophage subsets and fibrosis, we conducted Picro Sirius Red (PSR) staining over the immunostaining in a subset of the biopsies (n=6). This was done after carefully removing the cover slides, followed by capturing new images. Total $CD163^+$ macrophage area and the presence of S100A9/CD163 classical MDMs were semi-quantitatively analyzed with Fiji/ImageJ2 per staining [22]. TREM2, SPP1, and PLIN2 are highly expressed by tubuli and some biopsies showed minimal tubular CD163 uptake. This slight tubular CD163 staining could significantly skew quantitative colocalization analyses of sparse SPP1/CD163 macrophages and were therefore not performed. Thresholds of CD163 and S100A9 were manually determined in a selection of 4 biopsies and kept stable for samples in the same batch. Since the staining intensity between batches was not completely identical, thresholds could not be kept stable across batches and was again manually determined in a selection of 4 biopsies. To semi-quantify macrophage subsets, first, the outline of the kidney biopsy was automatically determined to specify the region of interest (ROI). To improve the accuracy of the semi-quantitative measurements in instances where edge or fold artifacts were detected along with the presence of the renal capsule or larger areas displaying non-specific staining, these regions were excluded from measurement process by exclusion from the ROI. Subsequently, macrophages expressing S100A9 were identified by determination of colocalization with CD163. Next, the % of the total biopsy area with dual-positive S100A9 and CD163 staining was calculated and compared between groups. Two NC biopsies demonstrated non-specific binding of CD163 and PLIN2 to erythrocytes, and one NC biopsy had excessive non-specific tubular CD163 staining leading to their exclusion from the overall CD163 measurement. However, since S100A9 staining was specific to immune cells, these particular biopsies were retained for subsequent analyses involving colocalization between CD163 and S100A9. In instances where data exhibited a log-normal distribution, a log transformation was applied, followed by a Welch ANOVA to assess disparities between AGN and NC, as well as

LN and NC and a T-test to assess differences between PR3-ANCA and MPO-ANCA patients. For datasets that did not conform to a normal distribution, non-parametric tests were utilized for comparative analysis. These analyses were performed using Graphpad Prism version 9.3.1.471 (GraphPad Software, La Jolla, CA, USA).

**Correlations between macrophage subtypes and histopathological/clinical features**

Paraffin sections of kidney biopsies were prepared and subjected to staining (hematoxylin and eosin (H&E), Periodic acid-Schiff (PAS) and Jones' silver methenamine staining) to perform a detailed histopathological examination by experienced nephropathologists (JR and JK). The total number of glomeruli, sclerosed glomeruli, cellular, fibrocellular and fibrous crescents, break of Bowman's capsule (BC), fibrinoid necrosis of glomerular capillaries (FN), acute tubular necrosis, percentage of interstitial fibrosis and tubular atrophy (IFTA), percentage of (tubulo-)interstitial inflammation (TIN), arteritis of larger arteries and arteriosclerosis were assessed. A chronicity score was calculated using the modified NIH chronicity index [15]. An activity index based on cellular and fibrocellular crescents, TIN, FN and BC break was used quantify histological disease activity. To look into correlations between markers of acute and long-term damage, we selected a limited set of variables (normal glomeruli, acute injury: modified activity index, % cellular crescents, %TIN; chronic injury: NIH chronicity score, estimated glomerular filtration rate (eGFR) at biopsy, and delta eGFR first year after biopsy) to avoid the need for multiple testing correction. eGFR was calculated using the CKD-EPI 2021 formula [23]. Spearman's Rho correlation tests were employed to assess the relationships between these variables and macrophage subsets. For histological comparative analysis between ANCA subtypes, in instances where data exhibited a normal distribution, t-tests were utilized, otherwise non-parametric tests were chosen.

**Supplementary Figures**

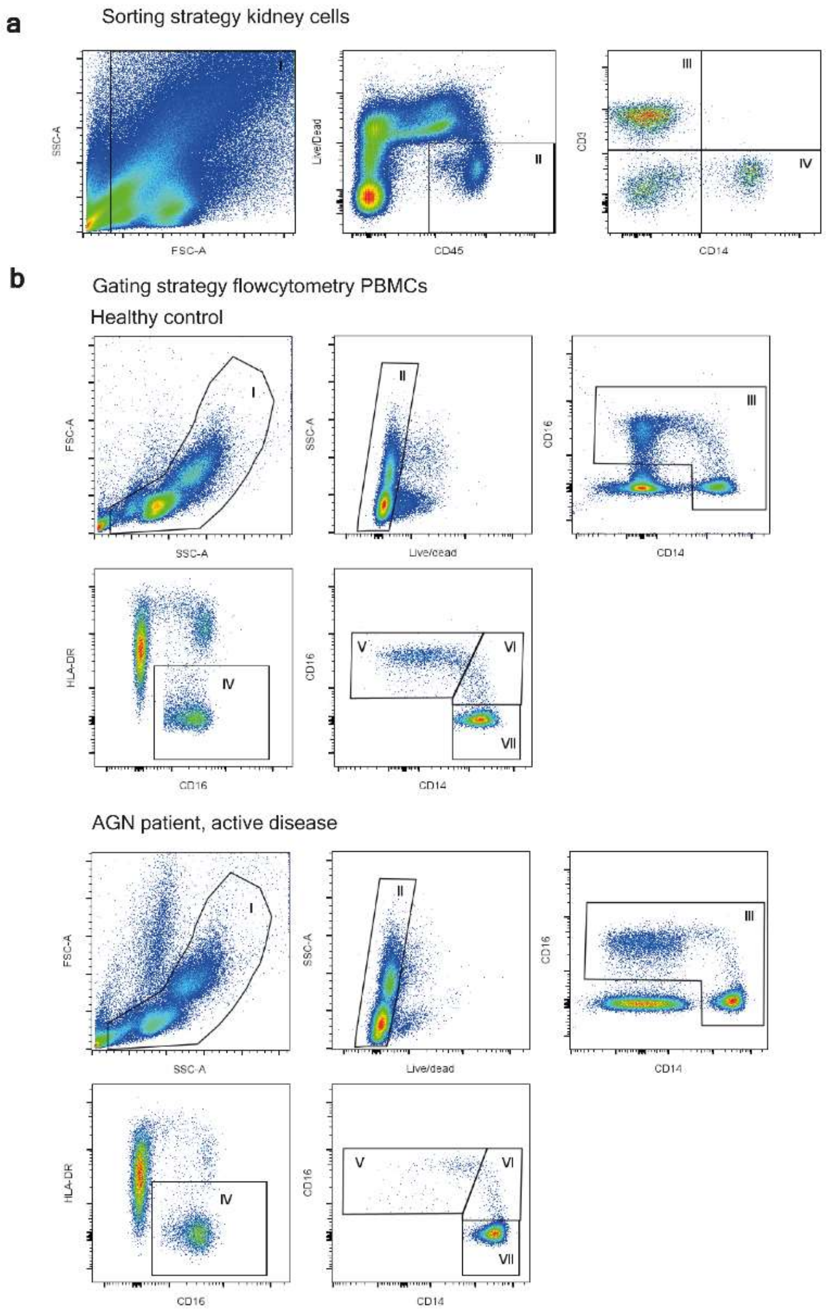

**Supplementary Fig. S1: Sorting and gating strategies**

**a**, Gating of kidney immune cell sort. FSC/SSC plot (gate I) was used to remove Residual RBC and debris. Gate I was presented on a CD45/Live/dead plot to sort living CD45+ immune cells (gate II). To estimate the number of T cells (gate III) and monocytes (gate IV), cells from gate II were plotted on a CD14/CD3 plot. **b**, Gating of monocyte subtypes, example of healthy control (up) and AGN patient (down). First, Residual RBC, debris and granulocytes were excluded on the FSC/SSC plot (gate I). Next, living cells were selected on the SSC-A/Live/dead plot (gate II) and presented on a CD14/CD16 plot to select $CD14^{+}$ and/or $CD16^{+}$ cells (gate III). HLA-DR- and CD16- cells were excluded (gate IV), followed by gating of monocyte subsets, $CD14^{++}/CD16^{-}$ (Classical) (gate VII), $CD14^{++}/CD16^{+}$ (intermediate) (gate VI), and $CD14^{+}/CD16^{++}$ (non-classical) (gate V) monocytes.

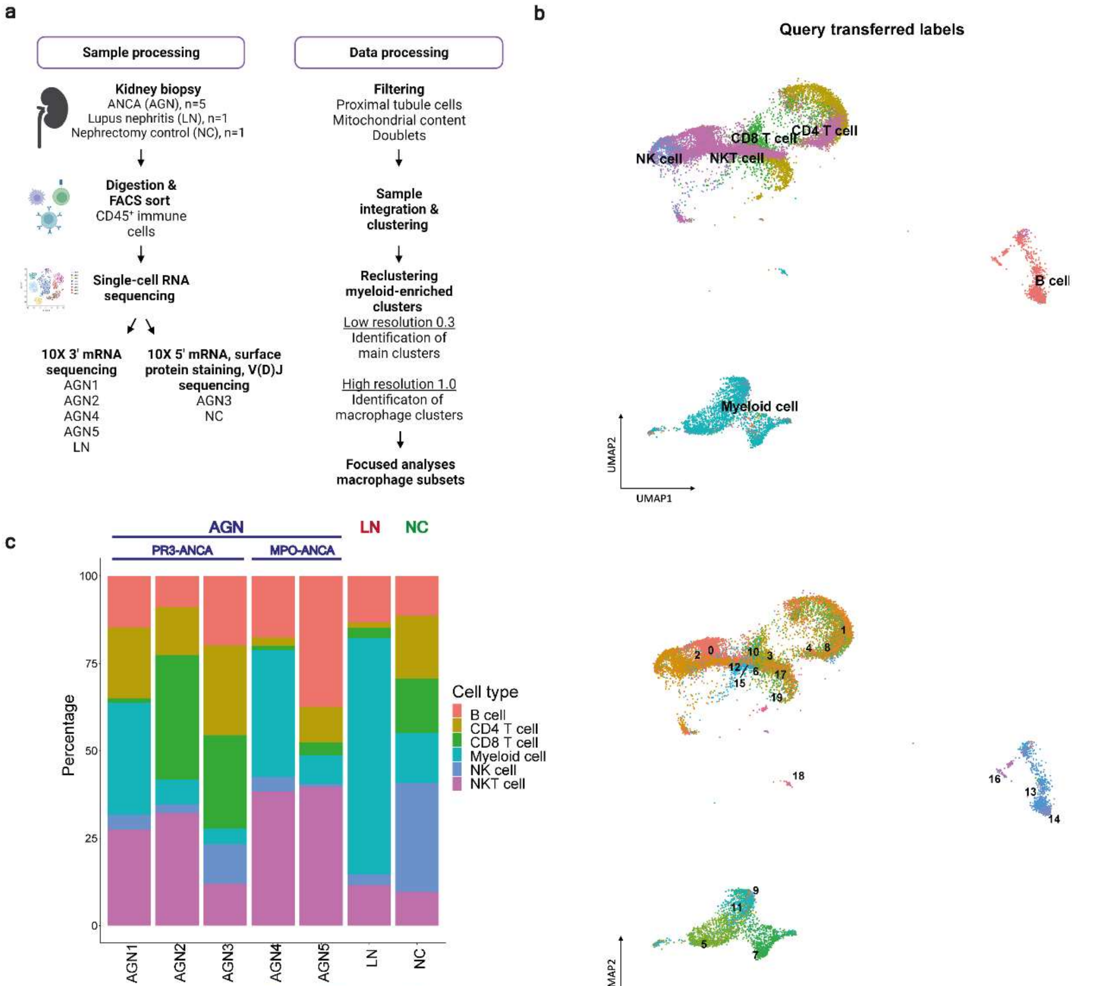

| Cell type | AGN | LN | NC |
|---|---|---|---|
| B cell | 20% | 13% | 11% |
| CD4 T cell | 14% | 2% | 18% |
| CD8 T cell | 14% | 3% | 16% |
| Myeloid cell | 18% | 68% | 14% |
| NK cell | 5% | 3% | 31% |
| NKT cell | 30% | 12% | 10% |

**Supplementary Fig. S2: Extended methods and total immune cell composition by mapping onto reference atlas**

**a**, Flowchart of sample and data processing. **b**, Uniform manifold approximation and projection (UMAP) visualizations of 25,485 single kidney immune cells acquired as in **a**, projected and annotated based on mature immune kidney cell atlas of Stewart *et al.* 2019. Upper panel, cells colored by cell type; lower panel, cells colored by cluster (**Fig. 1b**, left panel). **c**, Stacked-bar plot showing kidney immune cell type percentages per sample based on Stewart *et al* 2019 annotation. The table (lower panel) summarizes mean percentages per group.

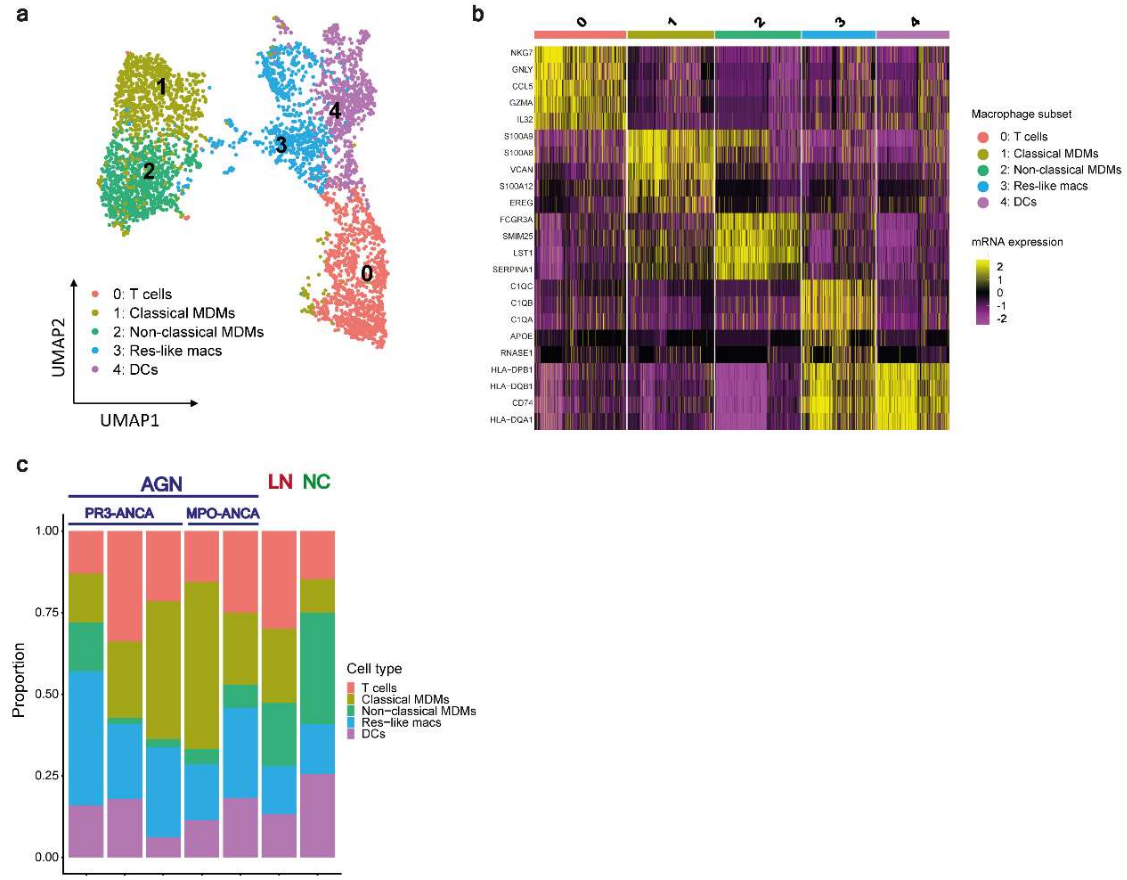

**Supplementary Fig. S3: Low-resolution subclustering identifies macrophages, T cells and dendritic cell clusters**

**a**, Uniform manifold approximation and projection (UMAP) visualization of sub-clustered myeloid-enriched clusters at low resolution (0.3). Five clusters were identified, containing 3 macrophage clusters. **b,** Heatmap showing scaled single-cell mRNA expression of top 5 markers. **c,** Stacked-bar plot of cell type proportions per sample. AGN, ANCA-associated glomerulonephritis; LN, lupus nephritis; NC, nephrectomy control; PR3, proteinase-3; MPO, myeloperoxidase; MDM, monocyte-derived macrophage; Res-like, resident like; DCs, Dendritic cells.

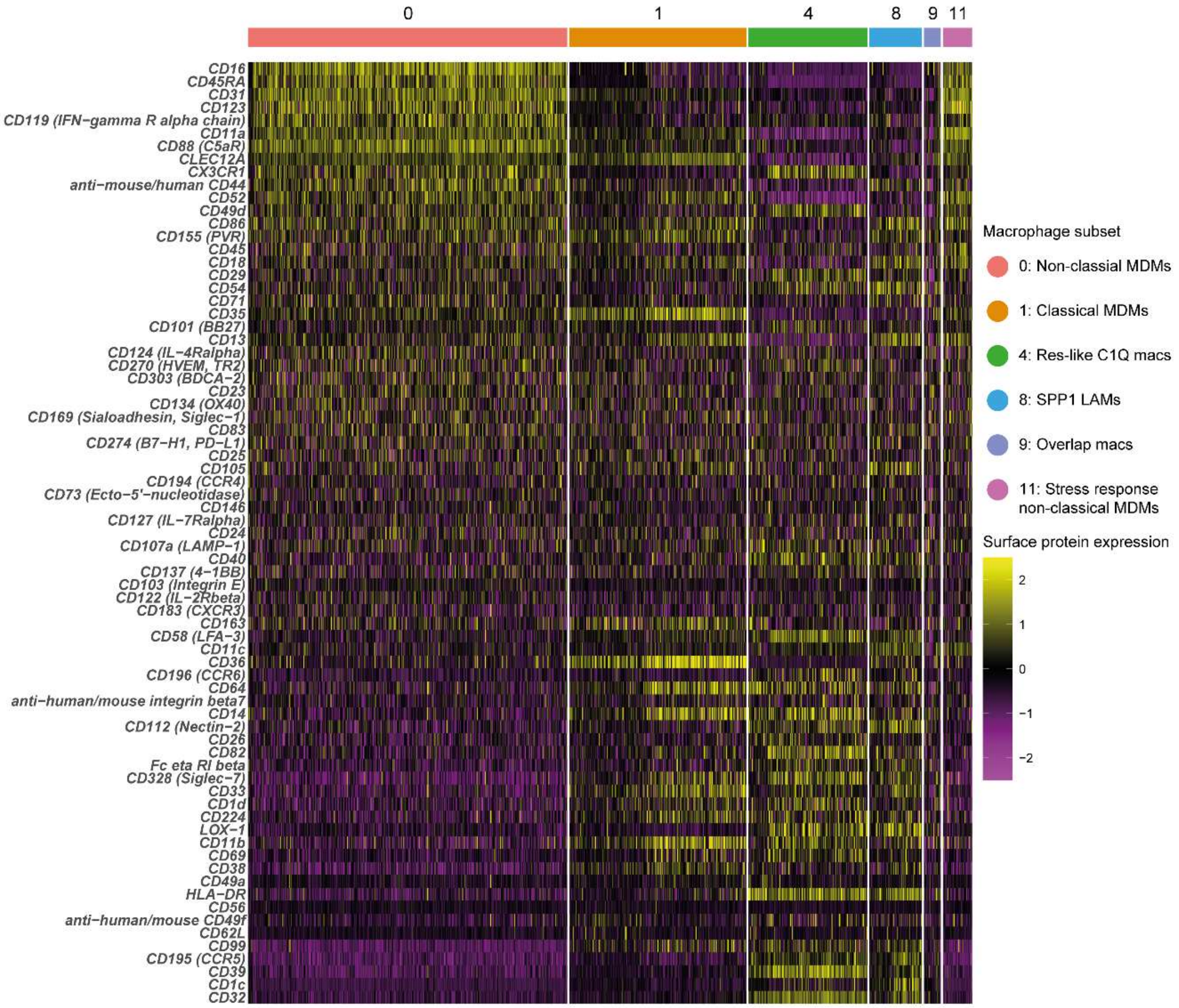

**Supplementary Fig. S4: Expression of monocyte/macrophage surface proteins per macrophage subset.** Heatmap showing scaled single-cell surface protein expression in two samples (AGN3 and NC). MDMs, monocyte-derived macrophage; Res-like *C1Q* macs, resident-like macrophages; LAMs, lipid-associated macrophages.

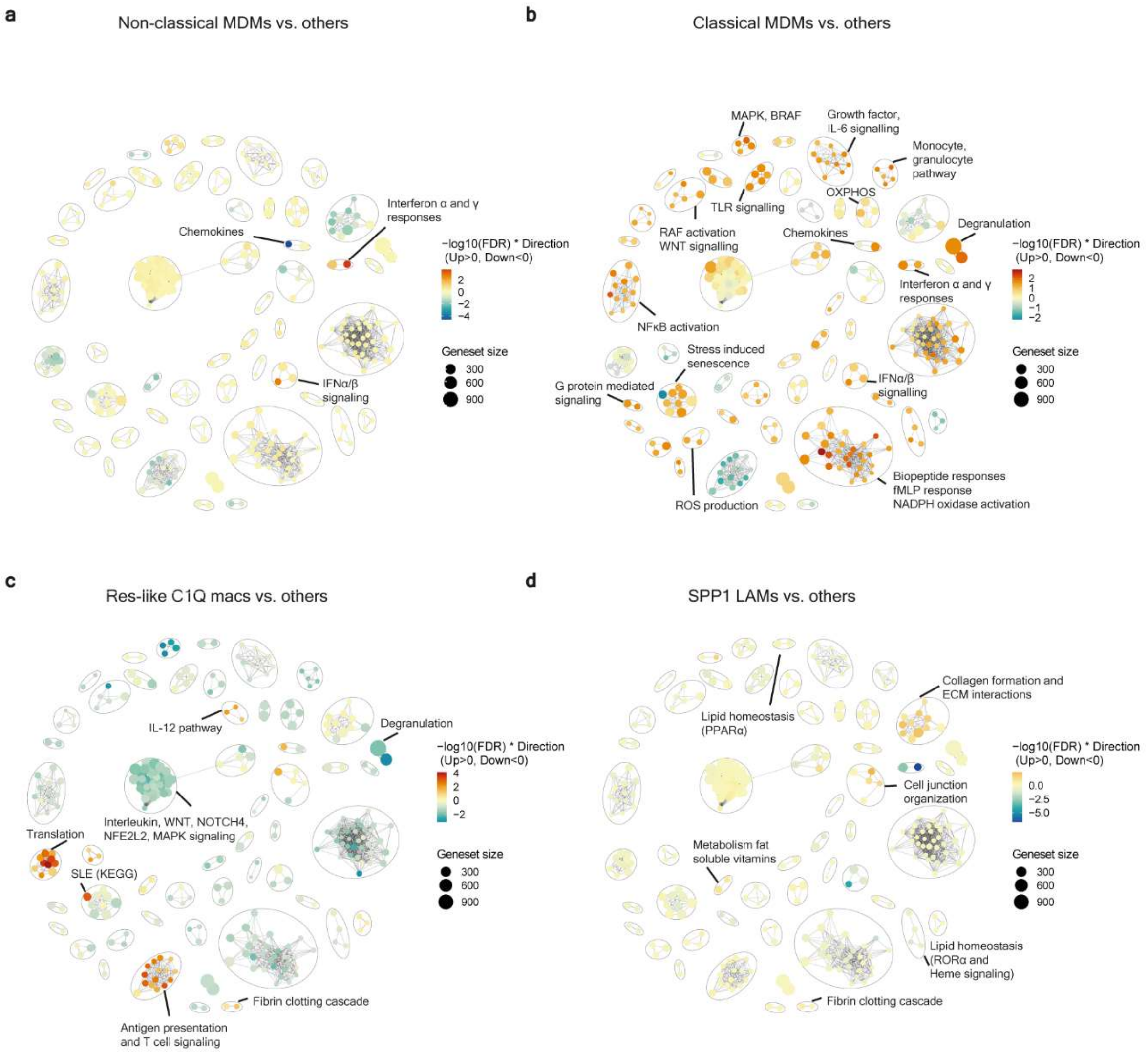

**Supplementary Fig. S5: GSEA shows enrichment of inflammatory pathways in classical MDMs and profibrotic genesets in *SPP1* LAMs. a-d,** Geneset enrichment analysis (GSEA) was performed comparing each of the main macrophage subsets with the remaining subsets using four MSigDB geneset collections (Hallmark, BioCarta, KEGG, Reactome). Enrichment networks are shown to group genesets based on genes they have in common. The nodes represent genesets that are significantly upregulated (red) or downregulated (blue) in the indicated macrophage subset. Overlap between genesets is represented by lines. Annotation was manually added. (**a**) Enrichment network for non-classical MDMs shows upregulation of interferon responses. (**b**) Classical MDMs express multiple genesets associated with MAPK signaling and inflammatory responses, such as degranulation, NFκB activation and toll-like receptor signaling. (**c**) Res-like *C1Q* macs express genesets associated with translation, classical complement activation and signaling of T cells. (**d**) *SPP1* LAMs show upregulation of genesets associated with collagen formation, elastic fiber formation, adipo-signaling. MDMs, monocyte-derived macrophage; Res-like *C1Q* macs, resident-like macrophages; LAMs, lipid-associated macrophages.

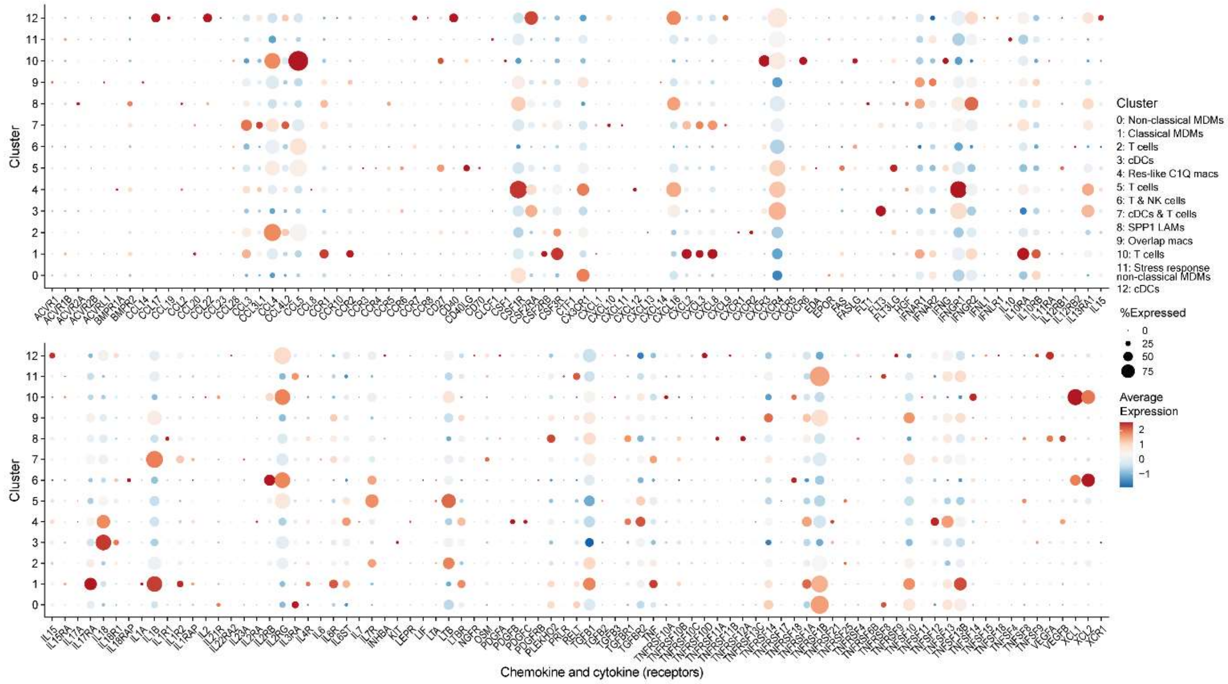

**Supplementary Fig. S6: Chemokine and cytokine (receptor) expression in all subclusters**

Dot plot illustrating chemokine and chemokine (receptor) expression in all 13 subclusters. MDMs, monocyte-derived macrophages; Res-like macs, resident-like macrophages; cDCs, classical dendritic cells; LAMs, lipid-associated macrophages; mac, macrophages

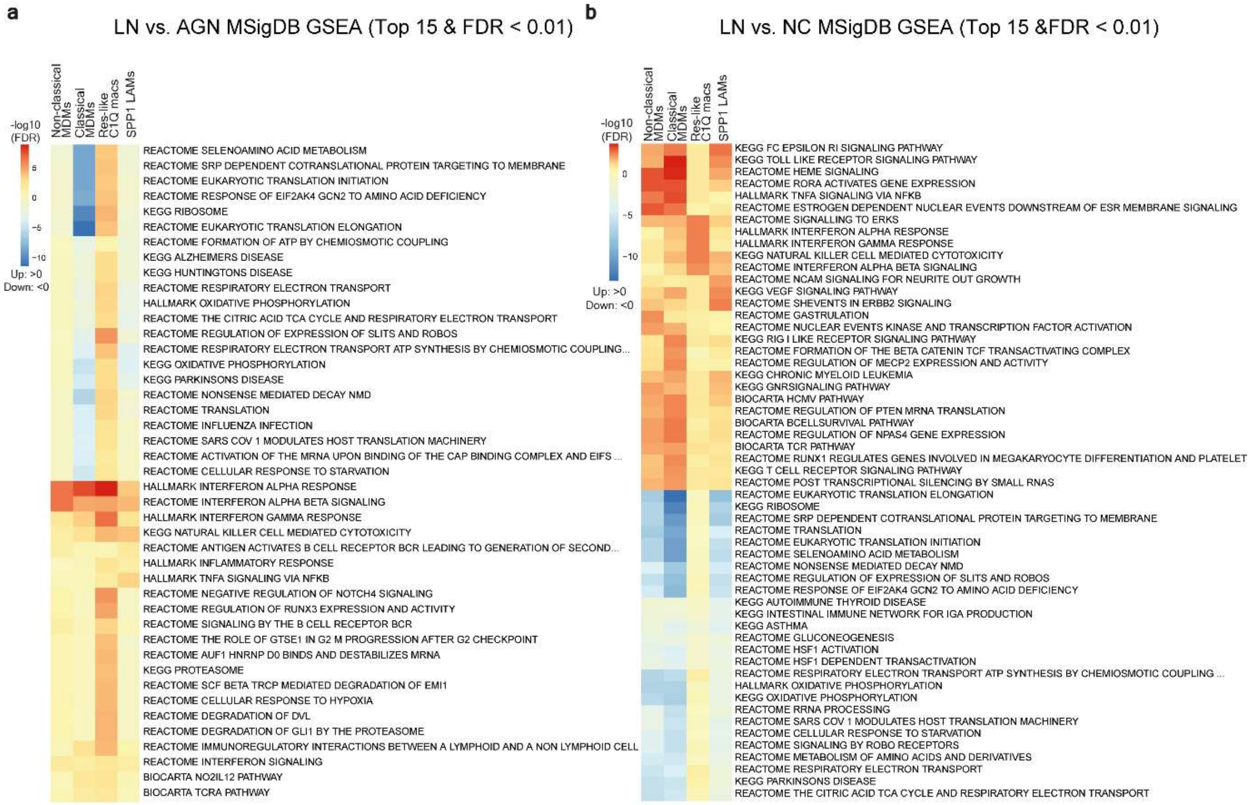

**Supplementary Fig. S7: LN disease-control macrophages show overall activation of interferon responses and specific activation of resident-like *C1Q* macrophages**

**a-b,** Geneset enrichment analysis (GSEA) of selected geneset collections to investigate disease-specific transcriptional changes within macrophage subsets. Heatmap of top 15 significantly (FDR < 0.01) upregulated (red) and downregulated (blue) genesets within macrophage subsets. (**a)** LN compared to AGN (**b)** LN compared to NC. C0, non-classical monocyte-derived macrophages (MDMs); C1, classical MDMs; C4, resident-like C1Q macrophages; C8, SPP1, lipid-associated macrophages; AGN, ANCA-associated glomerulonephritis; LN, lupus nephritis; NC, nephrectomy control.

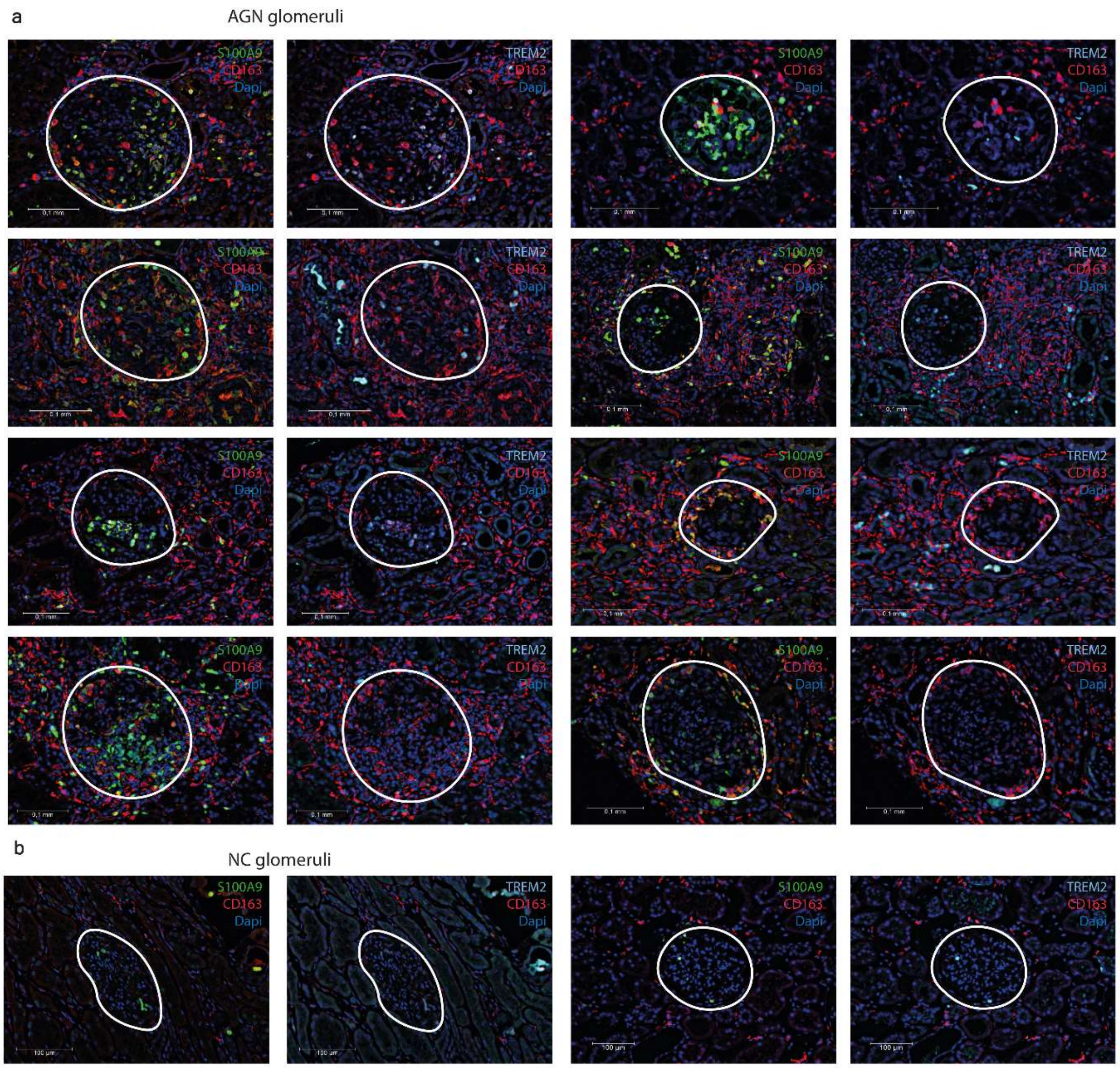

**Supplementary Fig. S8: Tubulointerstitial and glomerular infiltration of TREM2/CD163 macrophages is related to S100A9/CD163 macrophage accumulation in AGN**

**a-b,** Multi-color immunofluorescence stainings of kidney biopsies illustrating (left) S100A9/CD163 and (right) TREM2/CD163 expressing macrophages of (**a**) AGN patients (n=22) and (**b**) nephrectomy controls (n=8). AGN, ANCA-associated glomerulonephritis; NC, nephrectomy control.

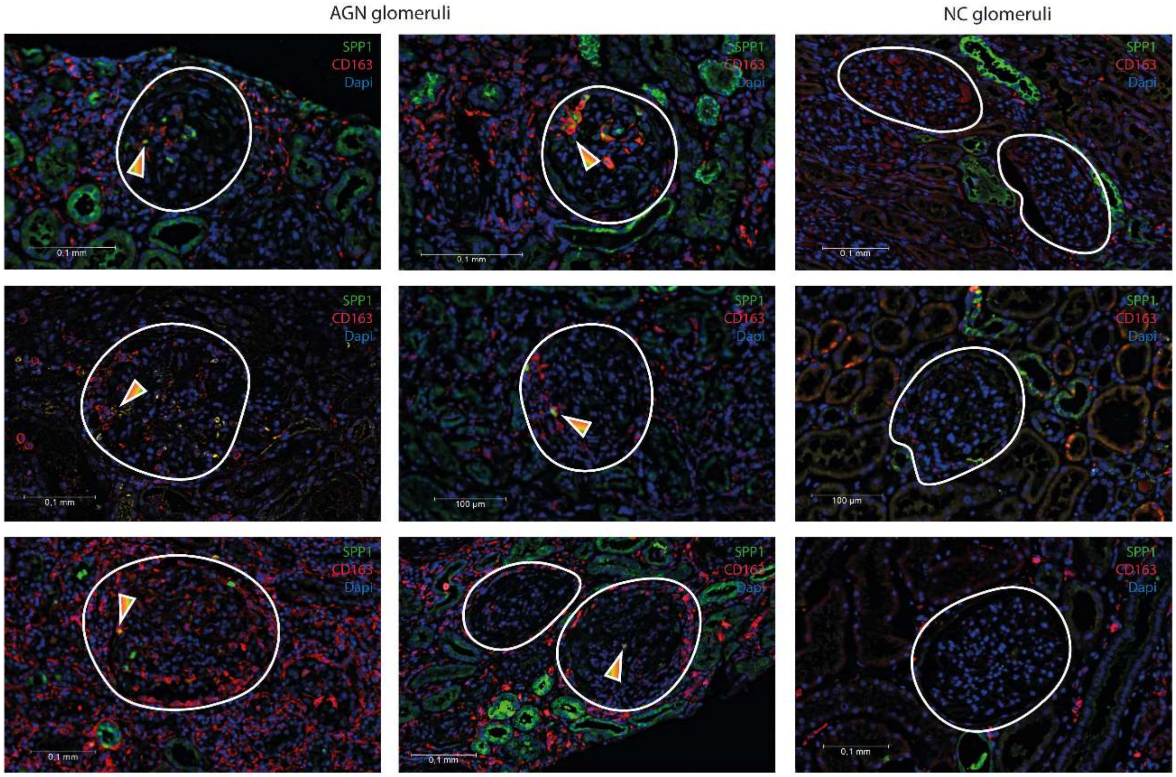

**Supplementary Fig. S9: SPP1/CD163 macrophages are located in glomeruli of AGN patients**
Multi-color immunofluorescence stainings of kidney biopsies illustrating SPP1/CD163 expressing macrophages (arrows) in kidney biopsies of AGN (n=22) and NC (n=8). Glomeruli are delineated by white circles. AGN, ANCA-associated glomerulonephritis; NC, nephrectomy control.

### Supplementary Tables

| | AGN1 | AGN2 | AGN3 | AGN4 | AGN5 | LN | NC |
|---|---|---|---|---|---|---|---|
| **Sex** | F | M | F | F | M | M | F |
| **Age, years** | 71 | 73 | 56 | 55 | 40 | 62 | 83 |
| **BMI** | 36 | 36 | 32 | 29 | 24 | 25 | 28 |
| **Serological subtype** | PR3 | PR3 | PR3 | MPO | MPO | | |
| **Disease activity score (BVASv3)** | 17 | 32 | 14 | 12 | 16 | | |
| **Damage score (VDI)** | 1 | 0 | 0 | 0 | 0 | | |
| **BSE, mm/U** | 33 | 29 | 81 | NA | NA | 32 | NA |
| **CRP, mg/L** | 14.6 | 58 | 30.1 | NA | 6.7 | 80.9 | 0.8 |
| **ANCA titer, kU/L** | 93 | 79 | 134 | 76 | 650 | | |
| **eGFR, mL/min/1,73m²** | 56 | 55 | 63 | 16 | 5 | 54 | 67 |
| **Immunosuppressive treatment** | MTX | MPNS | MPNS | MPNS, CYC, RTX | MPNS, PLEX | No | No |
| **Total MPNS dosage** | - | 2000mg | 2000mg | 3000mg | 2000mg | - | - |

**Supplementary Table S1: Patient characteristics kidney biopsies single-cell sequencing**

F, female; M, Male; PR3, proteinase-3; MPO, myeloperoxidase; BMI, body-mass index; BVASv3, Birmingham Vasculitis Activity Score version 3; VDI, Vasculitis Damage Index; eGFR, estimated Glomerular Filtration Rate; MTX, methotrexate; MPNS, methylprednisolon; CYC, cyclophosphamide; RTX, Rituximab; PLEX, Plasmapheresis; NA, not available; AGN, ANCA-associated glomerulonephritis; LN, lupus nephritis; NC, Nephrectomy Control;

| | AGN1 | AGN2 | AGN3 | AGN4 | AGN5 | LN | NC |
|---|---|---|---|---|---|---|---|
| **Total glomeruli, n** | 20 | 18 | 11 | 8 | 37 | 20 | 100 |
| **Sclerosed glomeruli, n (%)** | 6 (30) | 1 (6) | 3 (27) | 2 (25) | 23 (62) | 1 (5) | 11 (11) |
| **Crescent, n (%)** | 2 (10) | 8 (44) | 7 (64) | 5 (63) | 14 (38) | 1 (5) | 0 (0) |
| Cellular, n (%) | 2 (10) | 4 (22) | 7 (64) | 1 (13) | 2 (5) | 1 (100) | 0 (0) |
| Fibrocellular, n (%) | 0 (0) | 3 (17) | 0 (0) | 4 (50) | 2 (5) | 0 (0) | 0 (0) |
| Fibrous, n (%) | 0 (0) | 1 (6) | 0 (0) | 0 (0) | 10 (27) | 0 (0) | 0 (0) |
| **Mesangiocapillary glomerulonephritis** | - | + | - | - | + | - | - |
| **BC break** | - | active | - | fibrosed | fibrosed | - | - |
| **Fibrinoid necrosis** | + | + | + | + | + | + | - |
| **ATN** | - | + | + | - | + | - | - |
| **IFTA (%)** | 10 | 40 | 30 | 20 | 70 | 0 | 10 |
| **TIN (%)** | 20 | 60 | 20 | 20 | 20 | 0 | 10 |
| **Modified activity score** | 4 | 8 | 9 | 9 | 5 | 3 | 1 |
| **Modified chronicity score** | 4 | 6 | 6 | 4 | 11 | 1 | 3 |
| **Arteriitis larger arteries** | - | - | - | - | - | - | - |
| **Arteriosclerosis** | + | + | - | + | + | + | + |

**Supplementary Table S2: Histological findings kidney biopsies single-cell sequencing**.

ATN, Acute tubular necrosis; IFTA, Interstitial Fibrosis and Tubular Atrophy; TIN, Tubulointerstitial Nephritis; BC, Bowman's capsula; AGN, ANCA-associated glomerulonephritis; LN, lupus nephritis; NC, Nephrectomy Control. An activity score based on cellular and fibrocellular crescents, TIN, FN and BC break was used quantify histological disease activity. The chronicity score was calculated using the modified NIH chronicity index [14].

| Antibody | Clone | Fluorochrome | Manufacturer | Product number | Dilution |
|---|---|---|---|---|---|
| **Flowcytometry** | | | | | |
| Anti-HLA-DR | G46-6 | PercpC5.5 | BD | 560652 | 1:80 |
| Anti-CD14 | M5E2 | PEcy7 | Biolegend | 301813 | 1:80 |
| Anti-CD16 | 3G8 | APC-H7 | BD | 560195 | 1:25 |
| Live/dead | | eFluor506UV | eBioscience | 65-0866 | 1:50 |
| **Sort** | | | | | |
| Anti-CD3 | HIT3a | FITC | BD Pharmingen | 555339 | 1:3 |
| Anti-CD14 | 8G3 | PE | Sanquin | M1661 | 1:3 |
| Anti-CD45 | HI30 | APC-Fire750 | Biolegend | 982314 | 1:167 |
| Live/dead | | eFluor506UV | eBioscience | 65-0866 | 1:55 |
| **Immunofluorescence** | | | | | |
| Nuclear | | Hoechst | | | |
| Anti-PLIN2 | Guinea pig | | Progen | GP40 | 1:100 |
| Anti-Guinea pig | Donkey | FITC | Jackson | 705-095-148 | 1:100 |
| Anti-SPP1 | Rabbit polyclonal IgG | - | Abcam | ab8448 | 1:300 |
| Anti-S100A9 | Rabbit, IgG monoclonal JF096-8 | - | Bio-Connect | ET1702-73 | 1:100 |
| Anti-Rabbit | Goat | Alexa Fluor 633 | Invitrogen | A21070 | 1:500 |
| Anti-CD163 | Mouse IgG1 | - | ThermoFisher | MS-1103 | 1:250 |
| Anti-Mouse IgG1 | Goat | Alexa Fluor 568 | Abcam | ab175744 | 1:500 |
| Anti-TREM2 | Goat polyclonal IgG | - | Abcam | ab85851 | 1:50 |
| Anti-goat | Donkey | Alexa Fluor 750 | Abcam | ab175744 | 1:50 |

**Supplementary Table S3: Monoclonal antibodies for flowcytometry, cell sorting and immunofluorescence**

| | AGN1 | AGN2 | AGN3 | AGN4 | AGN5 | NC | LN | Total |
|---|---|---|---|---|---|---|---|---|
| **Sort** | | | | | | | | |
| Total cells sorted | 5,701,713 | 1,778,924 | 7,366,253 | 3,818,955 | 2,381,495 | 31,482,953 | 5,651,856 | 58,182,149 |
| Live, $CD45^+$ cells | 138,846 | 21,935 | 41,396 | 37,151 | 21,562 | 167,740 | 8,998 | 437,628 |
| %$CD14^+$ | NA | 3.2% | 2.9% | 15.2% | 5.5% | 7.8% | 18.3% | |
| %$CD3^+$ | NA | 75.5% | 46.3% | 56.0% | 57.8% | 57.8% | 38.5% | |
| **Sequencing** | | | | | | | | |
| Sequencing modality | 3’ | 3’ | 5’<br>- B and T cell receptor repertoires<br>- Surface proteins | 3’ | 3’ | 5’<br>- B and T cell receptor repertoires<br>- Surface proteins | 3’ | |
| Total cells after filtering steps | 466 | 2747 | 5553 | 1231 | 4583 | 8359 | 2546 | 25,485 |
| Cells myeloid-enriched clusters (% total) | 107 (23%) | 201 (7%) | 276 (5%) | 263 (21%) | 292 (6%) | 1,517 (18%) | 1,538 (60%) | 4,194 (17%) |

**Supplementary Table S4: Overview cell counts at selected sorting and data processing steps**
NA, not available; AGN, ANCA-associated glomerulonephritis; LN, lupus nephritis; NC, Nephrectomy Control.

| Annotation \ Cluster | 0 | 1 | 2 | 3 | 4 | 5 | 6 | 7 | 8 | 9 | 10 | 11 | 12 | 13 | 14 | 15 | 16 | 17 | 18 | 19 | Total cells (%) |
|---|---|---|---|---|---|---|---|---|---|---|---|---|---|---|---|---|---|---|---|---|---|
| B cell (n) | 0 | 2 | 0 | 1 | 0 | 3 | 0 | 42 | 0 | 2 | 0 | 18 | 1 | 594 | 627 | 8 | 97 | 0 | 0 | 0 | 1395 (6) |
| CD4 T cell (n) | 92 | 1764 | 1 | 320 | 1043 | 24 | 187 | 42 | 616 | 2 | 10 | 6 | 13 | 2 | 0 | 4 | 4 | 47 | 0 | 60 | 4237 (17) |
| CD8 T cell (n) | 51 | 5 | 1 | 937 | 110 | 9 | 272 | 3 | 108 | 0 | 371 | 5 | 38 | 0 | 0 | 10 | 3 | 0 | 0 | 0 | 1923 (8) |
| Myeloid cells (n) | 10 | 1 | 36 | 1 | 2 | 1179 | 0 | 1013 | 0 | 980 | 0 | 402 | 15 | 1 | 4 | 33 | 1 | 0 | 63 | 0 | 3741 (15) |
| NK cell (n) | 4 | 0 | 875 | 0 | 0 | 1 | 0 | 0 | 0 | 0 | 2 | 13 | 12 | 0 | 0 | 2 | 0 | 0 | 0 | 0 | 909 (4) |
| NKT cell (n) | 3730 | 1394 | 2165 | 1778 | 1221 | 65 | 711 | 21 | 391 | 0 | 588 | 364 | 575 | 51 | 12 | 142 | 9 | 62 | 0 | 1 | 13280 (52) |
| Total cells per cluster (n) | 3887 | 3166 | 3078 | 3037 | 2376 | 1281 | 1170 | 1121 | 1115 | 984 | 971 | 808 | 654 | 648 | 643 | 199 | 114 | 109 | 63 | 61 | 25485 (100) |
| % Of total myeloid cells | 0.3 | 0.0 | 1.0 | 0.0 | 0.1 | 31.5 | 0.0 | 27.1 | 0.0 | 26.2 | 0.0 | 10.7 | 0.4 | 0.0 | 0.1 | 0.9 | 0.0 | 0.0 | 1.7 | 0.0 | |

**Supplementary Table S5: Overview counts all immune cell clusters projected on mature immune kidney cell atlas Stewart *et al.* (2019).**
Myeloid cells contain the following groups: Mast cell, MNP-a/classical monocyte derived, MNP-b/non-classical monocyte derived, MNP-c/dendritic cell, MNP-d/Tissue macrophage, Neutrophil. MNP= mononuclear phagocyte. Grey columns indicate cell clusters containing ≥ 5% of total myeloid cells which were selected for reclustering.

| Sample / Cluster | AGN | | | | | LN | NC | Total cells |
|---|---|---|---|---|---|---|---|---|
| | AGN 1 | AGN2 | AGN3 | AGN4 | AGN5 | | | |
| **0** | 16 | 4 | 7 | 12 | 19 | 271 | 470 | 799 |
| **1** | 16 | 47 | 116 | 126 | 63 | 266 | 149 | 783 |
| **2** | 5 | 47 | 25 | 37 | 36 | 303 | 41 | 494 |
| **3** | 11 | 11 | 13 | 9 | 17 | 69 | 251 | 381 |
| **4** | 26 | 24 | 28 | 14 | 46 | 77 | 149 | 364 |
| **5** | 1 | 6 | 9 | 7 | 21 | 133 | 127 | 304 |
| **6** | 9 | 8 | 11 | 8 | 14 | 105 | 118 | 273 |
| **7** | 5 | 23 | 3 | 17 | 24 | 122 | 52 | 246 |
| **8** | 7 | 18 | 31 | 11 | 24 | 48 | 48 | 187 |
| **9** | 11 | 3 | 15 | 21 | 14 | 88 | 10 | 162 |
| **10** | 0 | 10 | 18 | 1 | 11 | 25 | 17 | 82 |
| **11** | 0 | 0 | 0 | 0 | 1 | 16 | 44 | 61 |
| **12** | 0 | 0 | 0 | 0 | 2 | 15 | 41 | 58 |
| **Total cells / sample** | 107 | 201 | 276 | 263 | 292 | 1538 | 1517 | 4194 |

**Supplementary Table S6: Overview counts per cluster and sample at high resolution (1.0) after reclustering myeloid-enriched clusters**

AGN, ANCA-associated glomerulonephritis; LN, lupus nephritis; NC, Nephrectomy Control.

| | ANCA (n=22) | SLE (n=6) | Healthy controls (n=8) |
|---|---|---|---|
| **Demographics** | | | |
| **Age at biopsy, years, mean (SD)** | 55 (20) | 34 (16) | 63 (15) |
| **Female sex, n (%)** | 9 (41) | 5 (83) | 2/7 (29) |
| **Disease characteristics** | | | |
| **ANCA subtype** | | | |
| PR3/MPO, n (%) | 11 (50) /11 (50) | - | - |
| **Use of immunosuppressive medication, n (%)** | 16 (73) | 3 (50) | - |
| Prednisone use, n (%) | 2 (9) | 3 (50) | - |
| Methylprednisolon <30 days, n (%) | 13 (57) | 0 (0) | - |
| Cyclophosphamide, n (%) | 5 (23) | 0 (0) | - |
| Rituximab < 1 year, n (%) | 2 (9) | 0 (0) | - |
| Plasmapheresis, n (%) | 2 (9) | 0 (0) | - |
| Methotrexate, n (%) | 1 (5) | 0 (0) | - |
| Mycophenolate mofetil, n (%) | 0 (0) | 2 (33) | - |
| Hydroxychloroquine, n (%) | 0 (0) | 3 (50) | - |
| **Laboratory time of biopsy** | | | |
| **eGFR, (CKD-EPI), ml/min/1,73 m2, median (IQR)** | 30 (39) | 57 (72) | - |
| **24h proteinuria, gram, median (IQR)** | 0.94 (4.93) | 5.32 | - |
| **ANCA titer, median (IQR)** | 134 (321) | - | - |
| **dsDNA titer, median (IQR)** | - | 1920 (2309) | - |
| **Biopsy findings** | | | |
| **Total glomeruli, n, median (IQR)** | 21 (21) | 16 (11) | 300 (360) |
| **Obliterated glomeruli, %, median (IQR)** | 14 (33) | 0 (35) | 1(9) |
| **Crescents** | | | |
| Cellular, %, median (IQR) | 8 (38) | 0 (5) | 0 (0) |
| Fibrocellular , %, median (IQR) | 21 (19) | 0 (0) | 0 (0) |
| Fibrous, % , median (IQR) | 0 (5) | 0 (0) | 0 (0) |
| **Break in Bowman's capsula, n (%)** | 14 (64) | 0 (0) | 0/8 (0) |
| **Presence of fibrinoid necrosis, n (%)** | 16 (73) | 1 (17) | 0 (0) |
| **Presence of acute tubular damage, n (%)** | 14 (64) | 1 (17) | 2 (25) |
| **IFTA, %, median (IQR)** | 20 (23) | 20 (43) | 0 (0) |
| **TIN, % median (IQR)** | 15 (13) | 15 (35) | 0 (0) |
| **Presence of arteriitis, n (%)** | 0/21 (0) | 0 (0) | 0 (0) |
| **Presence of arteriosclerosis, n (%)** | 11/18 (61) | 1/6 (17) | 4/7 (57) |
| **Activity score** | 6.0 (4.3) | 2.0 (1.8) | 0.0 (0.0) |
| **Chronicity score** | 4.0 (3.0) | 3.0 (6.0) | 0.5 (1.0) |
| **Lupus Nephritis class** | | | |
| II , n (%) | - | 1 (17) | - |
| IV + V, n (%) | - | 3 (50) | - |
| V, n (%) | - | 2 (33) | - |
| **CD163 staining, %, median (IQR)** | 3.0 (2.8) | 3.0 (1.8) | 0.62 (0.25) |
| **S100A9/CD163 staining, %, median (IQR)** | 0.32 (0.42) | 0.18 (0.32) | 0.07 (0.20) |

**Supplementary Table S7: Patient and histological characteristics extended cohort immunofluorescence stainings kidney biopsies.** SD, standard deviation; IQR, interquartile range; PR3, proteinase-3; MPO, myeloperoxidase; eGFR, estimated glomerular filtration rate; IFTA, interstitial fibrosis and tubular atrophy; TIN, tubulointerstitial nephritis; lupus nephritis class according to the International Society of Nephrology/Renal Pathology Society classification system [15].
